# Invasive plants affect native plant pollination through pollinator-mediated cross-boundary effects

**DOI:** 10.1101/2025.09.18.677159

**Authors:** Rebecca A. Nelson, Neal Williams, Fernanda Valdovinos, Susan Harrison

## Abstract

Invasive plants may facilitate or compete with native plants by changing the foraging behavior and/or abundance of shared pollinators. Such effects have been explored within single habitats; however, many pollinators are highly mobile and can cross ecological boundaries. Whether via shared pollinators, invasive plants affect native plants in neighboring habitats remains an open question. Here, we ask this question in a mosaic of 22 native-plant-dominated meadows on serpentine soil, embedded in a matrix of non-serpentine meadows. The non-serpentine meadows contain dense stands of two insect-pollinated invasive species: *Vicia villosa,* and *Centaurea solstitialis*. The serpentine meadows contain two common native species that co-flower and share pollinators with these invasives: *Trifolium fucatum* and *Helianthus exilis*, respectively. Across three years, we determined that the abundance ratio of each invasive species in the surrounding landscape to its co-flowering native species within focal patches was associated with decreased visitation rates, seed set, and functional importance in the plant-pollinator network for the native species. We conclude that invasive plants can have indirect, negative effects on native plants in neighboring habitats, via competition for shared pollinators.

**Open research statement:** All data and code are accessible to the public via: https://figshare.com/projects/Data_for_Invasive_plants_affect_native_plant_pollination_through_pollinator-mediated_cross-boundary_effects/243587

## Introduction

Cross-boundary spillover occurs when organisms move from one habitat or ecosystem to a neighboring one (Polis et al. 1997, Tscharntke et al. 2005, Scherer-Lorenzen et al. 2022). Spillover can drive community assembly, maintain biodiversity, shape how habitats respond to anthropogenic global changes (Scherer-Lorenzen et al. 2022, Peller and Altermatt 2024), and support ecosystem functions and services, such as nutrient cycling and pollination (Polis et al. 1997, Tscharntke et al. 2005, Scherer-Lorenzen et al. 2022).

As highly mobile organisms, pollinators often forage across habitat boundaries and rely on spatially heterogeneous landscapes for floral resources (Hackett et al. 2024). Spillover of pollinators has been well studied in croplands, which provide tractable systems to explore spillover mechanisms and their potential consequences for crop pollination services (Tscharntke et al. 2005, Winfree et al. 2007, Ekroos et al. 2008). Proximity of fields to wildland habitats, for example, can enhance crop visitation and pollination through the spillover of wildland pollinators into croplands (Kremen et al. 2002, 2004, Ekroos et al. 2008, Williams 2011, Garibaldi et al. 2013).

Whether findings from cropland-wildland boundaries are applicable to wildland-wildland boundaries, which tend to be more diverse and spatially complex, remains an open question (Ponisio et al. 2016, Hackett et al. 2024). Although some evidence suggests that spillover can increase pollinator diversity and native plant fitness at wildland-wildland boundaries (Artz and Waddington 2006), other evidence suggests that bee movement between habitat patches is rare despite landscape heterogeneity (Harmon-Threatt and Anderson 2023).

Within a single habitat, plant species can have indirect negative or positive effects on each other’s reproduction via competition or facilitation for shared pollinators (Brown et al. 2002, Braun and Lortie 2019). Via competition for shared pollinators, invasive plant species can decrease native plant pollinator visitation, increase heterospecific pollen transfer, decrease native plant fitness (Brown et al. 2002, Morales and Aizen 2002, Traveset and Richardson 2006, Parra-Tabla et al. 2021) and/or increase the functional importance of the invasive plant species in the plant-pollinator network (Valdovinos et al. 2009, Parra-Tabla et al. 2021, Dritz et al. 2023). Invasive plant species can also have facilitative effects on native plant species by acting as “magnet” species that attract more pollinators to the area, increasing native plant pollinator visitation and fitness (Rodriguez 2006, Braun and Lortie 2019, Etter et al. 2022). Whether invasive plants can have indirect effects on native plant reproduction via pollinator spillover across habitats remains to be determined. If invasive plants decrease native plant fitness via spillover competition for shared pollinators, then habitats previously thought to be spatial refugia from the impacts of invasive species (e.g. serpentine grasslands, (Harrison et al. 2000) are not necessarily safe from their pollination-mediated negative effects (Harrison et al. 2000). Exploring this issue has implications for the management of refugia habitats that contain rare native plants (Case et al. 2016).

California serpentine grasslands are considered spatial refugia for native plants because their low-nutrient serpentine soils, high in heavy metals, prevent most invasive plant species from establishing in contrast to the highly invaded, non-serpentine soils with which they co-occur (Harrison et al. 2006, Harrison and Rajakaruna 2019). Serpentine-non-serpentine habitat boundaries allow us to investigate whether invasive, non-serpentine plant species can have cross-boundary effects on native serpentine plant pollination via pollinator spillover. The patchy nature of serpentine-non-serpentine mosaics allows for spatially replicated data.

In a three-year observational study, we examined whether the abundance ratios of two invasive species in non-serpentine grasslands to their co-flowering native species in adjacent serpentine grasslands were associated with decreased pollinator visitation rates, seed set, and contribution to plant-pollinator network structure for the native species. We hypothesized that as the ratio of invasive flowers to native flowers around the boundary increased, pollinator visitation to native plants would (1a) decrease in quantity, (1b) change in species composition, and (1c) decrease in quality; (2) native plant seed set would decrease; and (3) the functional roles of the native species in these plant-pollinator networks would decrease while those of the invasive species would increase.

## Methods

### Study System

This study took place at the University of California McLaughlin Reserve in the North Inner Coast Range of California (Lake County, CA 38.861°N, 122.408°W), wherein a mosaic of native wildflower-dominated serpentine grasslands and invaded non-serpentine grasslands occur within a patchy landscape (Harrison 1999). The area has a Mediterranean climate with hot, dry summers and cool, wet winters (Harrison et al. 2000).

We focused on two pairs of taxonomically related and co-flowering plant species; within each pair, one species was an invasive species growing on non-serpentine (sandstone-derived) soils and the other was a native species growing on serpentine soils. The invasive hairy vetch (*Vicia villosa* Roth, Fabaceae) co-flowers with the native bull clover (*Trifolium fucatum* Lindl., Fabaceae) in spring (April-May) (Baldwin and Goldman 2012, Harmon Threatt and Kremen 2015). The invasive yellow star-thistle (*Centaurea solstitialis* L., Asteraceae) co-flowers with the serpentine sunflower (*Helianthus exilis* A. Gray, Asteraceae) in summer (July) (Wolf et al. 1999, Barthell et al. 2001, Baldwin and Goldman 2012).

### Study Design

We conducted a three-year observational study of 22 naturally occurring meadows that contained serpentine-non-serpentine habitat boundaries. A multi-scale approach allowed for multiple units of replication across increasing biological scales: (1) the scale of an individual flowerhead, (2) the scale of a focal patch of native plants, defined as a contiguous, conspecific cluster of plants that co-flowered, (3) the scale of a meadow, an area of contiguous grassland containing a serpentine-non-serpentine boundary as defined by a geologic contact between soils, averaging ∼ 250 m in radius.

We selected focal meadows based on the following criteria: (1) contained a serpentine-non-serpentine habitat boundary, (2) had the focal native plant species flowering, and (3) was located at least 100 m away from any other meadow sampled. The total sampling region across all meadows sampled spanned an area of approximately 28.33 square kilometers with meadows separated from their nearest neighboring meadow by 100-300 m. Within our meadows, we observed plant-pollinator interactions at all patches of the focal native plants within a given meadow. Across all years, we observed pollinator visitation on a total of 50 native clover patches for spring and 44 native sunflower patches for summer. Within focal patches, we randomly selected and tagged individual flowerheads of focal native plants as follows: each native plant flowerhead had to be (1) located within a focal patch and meadow, (2) freshly opened with no signs of florivory, and (3) on different individuals of each focal native plant. We monitored a total of 850 individual clover flowerheads in 2024 and 700 individual sunflower flowerheads in 2023 and 2024, tagging 25-50 individual flowerheads per patch.

### Floral Abundance and Visitation to Focal Plants

We estimated the abundance of inflorescences as floral units within each focal patch using a log-binned method (10^1^ – 10^3^; (Mola and Williams 2018). We defined a floral unit as a single flowerhead or part of a multiple head from which a medium-sized bee would have to fly rather than walk to reach another unit of the same species (Lopezaraiza–Mikel et al. 2007).

We measured the number of pollinator visits to focal native plant patches and individual flowerheads within patches, only during calm and sunny conditions when focal invasive and native flower species were co-blooming. We considered a floral visit to have occurred if a floral visitor contacted the reproductive parts of the flower for at least one second. We visually observed and recorded all floral visitors to individual flowerheads within the patch and/or individually tagged flowerheads during three-minute observation periods. We surveyed spring pollinators on 3/9/-4/29/2022, 4/20-5/27/2023, 4/154-5/11/2024 and surveyed summer pollinators on 7/3-7/31/2022, 7/5-8/2/2023, and 7/8-7/31/2024, sampling each patch at least six times (approximately 1.5 observer hours patch^-1^year^-1^).

### Species and Morphospecies Identities

We photographed and/or netted voucher specimens of floral visitor morphospecies, which were identified by experts at the Bohart Museum of Entomology (bees, T. Zavortink; flies, S. Letana) supplemented by our use of iNaturalist and field guides (see Table S3). We sorted floral visitors into morphospecies based on matching observed visitors to our voucher specimens size, color, and shape, consistent with the methods of other network studies (DeSousa et al. 2025, Watts et al. 2016, Simonoak and Burkle 2014). For most bees, wasp and butterfly visitors, we identified morphospecies at the species or genus level, while for moths, flies, and beetle visitors, we identified most morphospecies at the genus and family level. In cases of multiple distinct morphospecies within a taxon, we noted the color or size differences used to distinguish them (Table S3). All poorly resolved taxa (unable to be identified beyond bee, fly or moth) were excluded from analyses. We paused timed observations when collecting insects and resumed timing once insects were collected. We aimed to collect insects once they left the focal area to minimize effects on visitation data. All voucher specimens were retained at the Michigan State Kellogg Biological Station’s Entomological Collections.

The only prevalent non-native pollinator in our system was the Eurasian honeybee (*Apis mellifera*), which in our study area was feral as opposed to coming from human-managed hives. Because honeybees are highly abundant visitors to invasive plants such as starthistle and vetch (Barthell et al. 2001), and because they may exert significant negative effects on native pollinators (Thomson 2004, Page and Williams 2023a, b, Travis et al. in press), we considered them both separately and combined with other pollinator species in the analyses described below.

### Visit Quality for Focal Plants

To measure heterospecific pollen transfer, in spring of 2022 and summer of 2024, we collected focal native plant stigmas from 6 clover patches in 6 meadows and from 6 sunflower patches in 5 meadows. We randomly selected 12 stigmas per patch from 12 individual plants. Stigmas were preserved in 70% ethanol and then scored using a light microscope. Prior to scoring, we mounted stigmas in fuchsin-tinted glycerin gel on microscope slides. Clover stigmas were directly mounted, while sunflower stigmas were softened with 0.1 M KOH for two hours at 60 °C and rinsed with deionized water before mounting to improve pollen visibility. For each stigma, we counted the number of conspecific and heterospecific pollen grains directly adhered to the surface and the number within a 5-conspecific pollen grain diameter of the stigmatic surface, since grains may have been dislodged during mounting.

### Focal Plant Seed Set

We measured seed set for focal native plants to characterize plant fitness. Our focal native plant species are annuals. We bagged all monitored flowerheads as seeds were maturing to deter seed predators and collected seedheads as they were senescing. We counted seeds per pod and weighed total seed mass for clovers, and counted seeds per flowerhead for sunflowers. Seed set data for spring were from 2022 and 2024, and for summer from 2023 and 2024.

### Meadow Plant-Pollinator Networks

We surveyed plant-pollinator networks across all meadows. Each meadow was surveyed at least 6 times per year for a total of approximately 6 observer hours meadow^-1^year^-1^). In each meadow, we placed a 100 m transect that avoided meadow edges by greater than 10 m. Each sample round, we walked each transect once, recording all flower visiting insects within 1 m for the transect and the identity of the flowering forb species which they visited. At the same time, we recorded the identity and flower abundance of all flowering plants within 1 m of the same transect (Lopezaraiza–Mikel et al. 2007). We sorted taxa into morphospecies using the approach described above.

### Floral Neighborhoods

We calculated the mean ratio of invasive to native plant floral abundance. Counts of invasive species were made within a 250 m radius of the focal native plant patch. The mean ratios were calculated on repeated visits across the flowering season and natural log-transformed. We use 250 m as our focal scale because this is the typical radius of a meadow (Moore et al. 2011) and is also a scale at which pollinators forage within a given meadow (Alignier et al. 2023). We also made counts at two larger radii (500 and 1000 m) to check for consistency of results across scales. We acknowledge that at these larger distances flower counts of the invasive species are not fully independent and so do not use them as our primary analysis.

### Data Analysis

To test (H1a) the effects of invasive to native plant ratio on the quantity and quality of visitation to native plants decreased with an increasing ratio of invasive to native plant abundance, we fit separate negative binomial generalized mixed effects models for each season (spring and summer) at both the individual flowerhead (see Supplemental Results) and patch scales, testing for a fixed effect of invasive-to-native-plant ratio and a random effect of meadow-year on the following pollinator response variables: pollinator morphospecies richness, the number of pollinator visits, number of western honeybee (*Apis mellifera*) visits, number of non-honeybee pollinator visits to the native plant species, and (H1c) the total number of invader heterospecific pollen grains present on native plant stigmas. The meadow-year random effect controlled for variation in years as well as spatial variation among meadows not related to invasive to native plant ratio.

To test (H1b) the effects of invasive to native plant ratio on pollinator composition on focal native plants, we ran PERMANOVAs with a Bray Curtis Dissimilarity Index using the Adonis2 function in the vegan R package (Dixon 2003) and analyzed generalized linear models of multivariate pollinator abundance using the mvabund package (Wang et al. 2012).

To test (H2) the effects of invasive to native plant ratio on native plant seed set, we fit mixed effects models, testing for a fixed effect of invasive-to-native-plant ratio and a random effect of meadow-year on seed set response variables (total number of seeds per flowerhead for both species and mean seed mass for clovers) for each season (spring and summer). We fit negative binomial models for sunflower seed counts and linear models for clover seed masses and counts.

To test (H3) the effects of invasive to native plant ratio on the functional importances of native and invasive plants in their plant-pollinator networks, we fit linear models to test for a fixed effect of mean invasive to native plant ratio at the meadow scale on network properties using the bipartite and igraph packages (Dormann et al. 2009, Csardi et al. 2025) (see Supplement). We examined 3 network metrics. (1) Individual species contribution to nestedness, which measures how strongly our focal plant species influence the overall structure of the network by attracting both generalist and specialist pollinators (Dormann et al. 2009, Csardi et al. 2025). (2) The Müller index, which measures perceived apparent competition for shared pollinators among focal plant species (Müller et al. 1999, Bergamo et al. 2021, Page and Williams 2023a), and allows us to examine whether the native or invasive plants receive more of the shared pooled of pollinator visits within the network. (3) Jaccard’s and Sorensen’s similarity in pollinators between focal invasive-native plant pairs: (Jaccard 1908, Sorensen 1948) averaged across sampling periods to examine whether invasive to native plant ratio correlated with how much focal plant species shared pollinators.

All data analyses were performed in R version 4.4.1 (R Core Team 2024) using the lme4 MASS R, and glmmTMB packages (Bates et al. 2009, Brooks et al. 2017).

## Results

In total, we observed 2309 individual pollinator visits to native clover patches, 138 visits to individual clover heads, 6021 individual pollinator visits to native sunflower patches and 1703 visits to individual sunflower heads. We observed a total of 11858 pollinator visits across all spring plant-pollinator networks including 134 unique floral visitor morphospecies (64 of which visited clovers) and 91 unique plant species. We observed a total of 11355 pollinator visits across all summer plant-pollinator networks including 119 unique floral visitor morphospecies (83 of which visited sunflowers) and 42 unique plant species (see Supplement). In spring, the log-transformed mean invasive vetch to native clover floral abundance ratio varied from -1.13 to 7.12 at 250 m, from -0.74 to 7.33 at 500 m, and 0.0025 to 8.95 at 1000 m. In summer, the log-transformed mean invasive thistle to native sunflower floral abundance ratio varied from -3.04 to 7.06 at 250m, from -3.04 to 7.06 at 500m, and from 0.21 to 7.41 at 1000 m.

### Pollinator visitation to Native Plants

Consistent with H1a, the quantity of pollinator visitation at the patch scale varied with boundary context in both spring and summer. In spring, pollinator morphospecies richness and total number of pollinator visits on the native clover decreased with increasing invasive-to-native-plant ratio (hereafter referred to as I:N ratio) (Figure 1A & B, Table 1), as did the total numbers of honeybee (Table 1), and non-honeybee visitors (Table 1).

**Figure 1.**
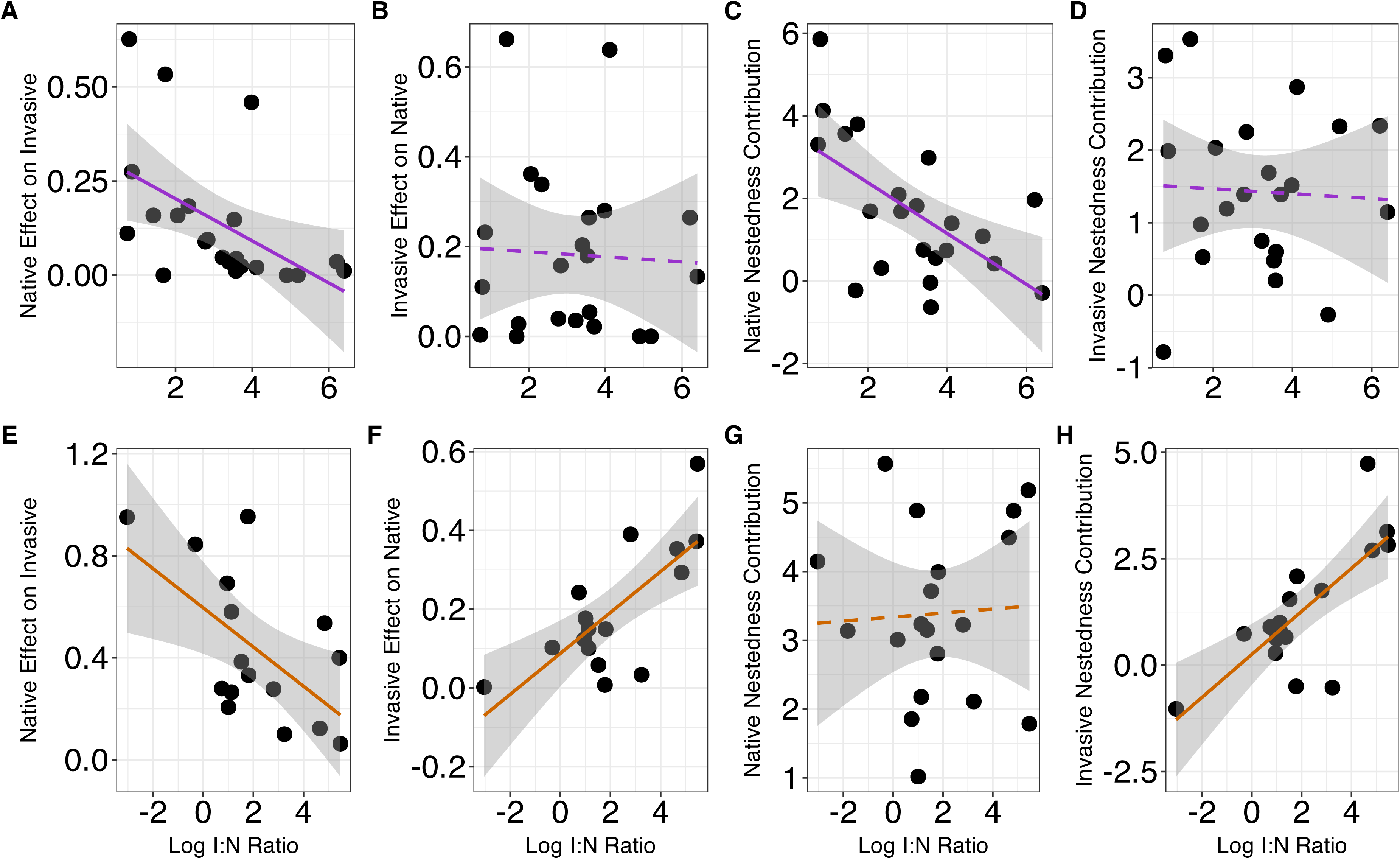
The relationship between invasive plant floral abundance at 250 M to native plant floral abundance for (A) pollinator morphospecies richness and (B) number of pollinator visits, and (C) mean seed mass per seedpod measured in grams for bull clover in the spring and for (D) pollinator morphospecies richness and (E) number of pollinator visits, and (F) total number of seeds per floral unit for serpentine sunflower in summer. The invasive plant for spring is hairy vetch, and the invasive plant for summer is yellow star-thistle. Each dot represents the mean values for a given patch over a given season. Solid regression lines denote a statistically significant relationship at an alpha level of 0.05.

**Table 1.**
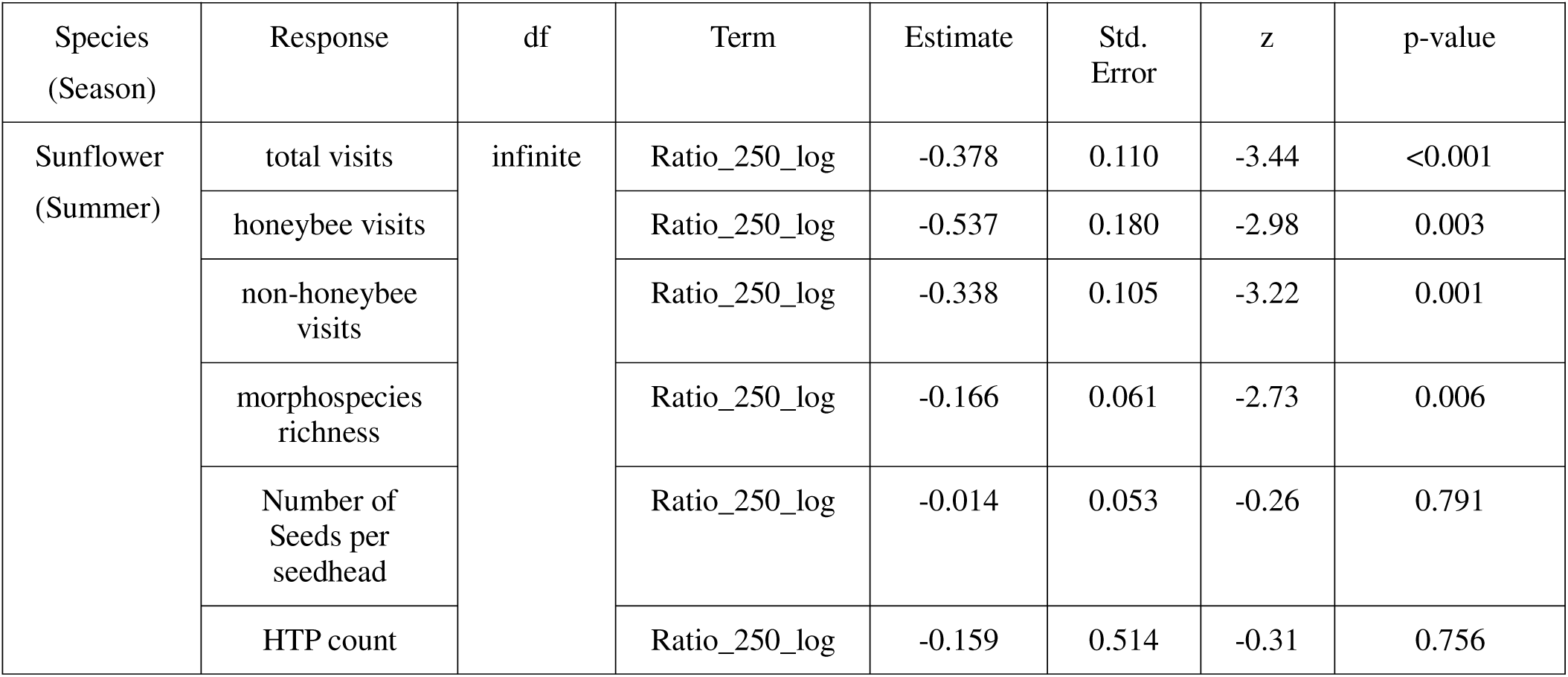
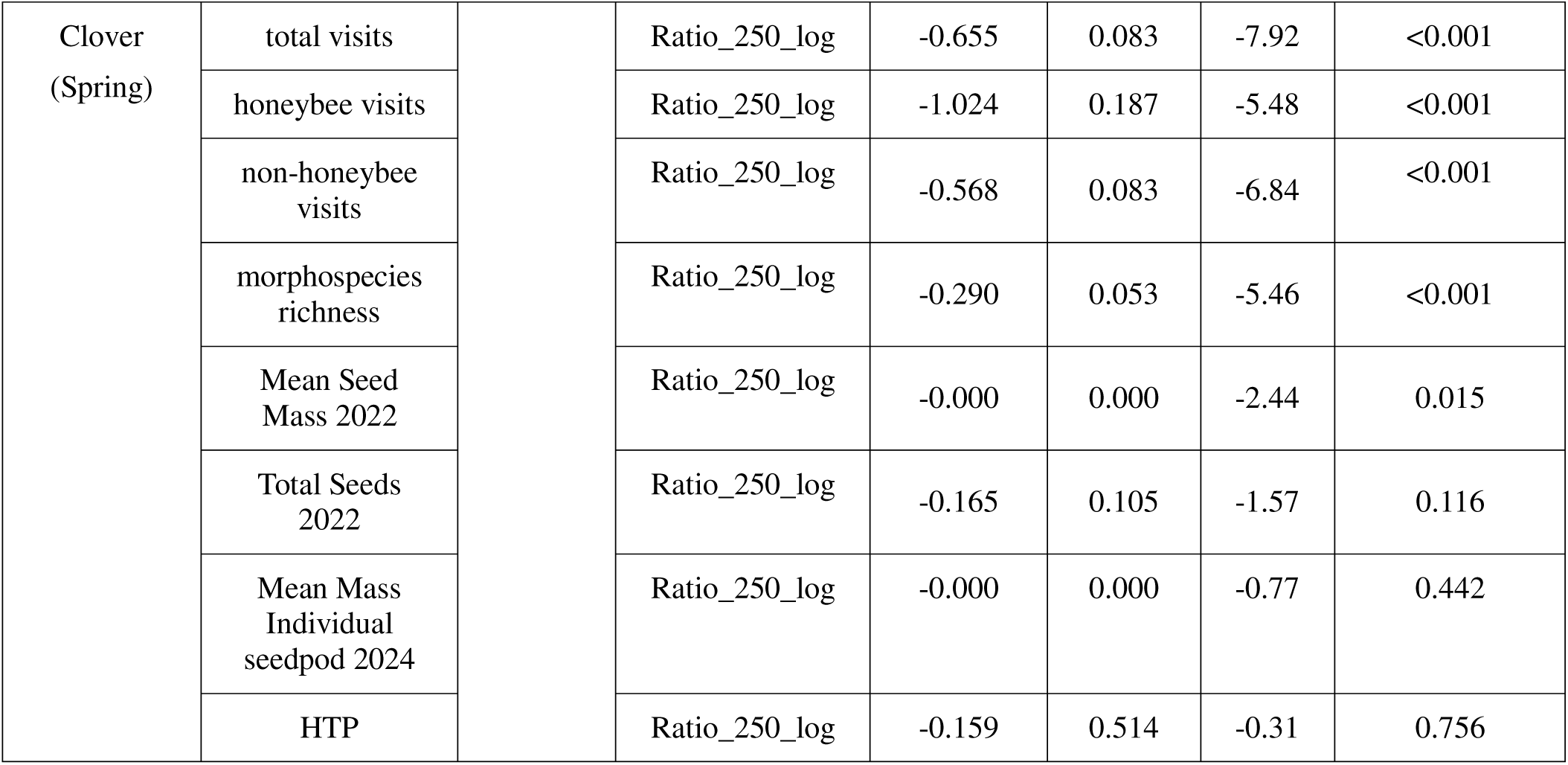
Pollinator Responses by Boundary Context. The statistical outputs of negative binomial and Gaussian for clover seed set mixed effects models, testing for an effect on log-transformed 250 m invasive to native plant ratio on patch-level visitation, seed set, and heterospecific counts for both spring native bull clovers (*T. fucatum*) and summer native serpentine sunflowers (*H. exilis*). All models had a random of effect of unique meadow-year. Because we used the glmmTMB package, all models had a test statistic of z and thus infinite degrees of freedom associated with the z-distribution. The alpha level for significance was 0.05.

| Species<br>(Season) | Response | df | Term | Estimate | Std.<br>Error | z | p-value |
| --- | --- | --- | --- | --- | --- | --- | --- |
| Sunflower<br>(Summer) | total visits | infinite | Ratio_250_log | -0.378 | 0.110 | -3.44 | <0.001 |
|  | honeybee visits |  | Ratio_250_log | -0.537 | 0.180 | -2.98 | 0.003 |
|  | non-honeybee<br>visits |  | Ratio_250_log | -0.338 | 0.105 | -3.22 | 0.001 |
|  | morphospecies<br>richness |  | Ratio_250_log | -0.166 | 0.061 | -2.73 | 0.006 |
|  | Number of<br>Seeds per<br>seedhead |  | Ratio_250_log | -0.014 | 0.053 | -0.26 | 0.791 |
|  | HTP count |  | Ratio_250_log | -0.159 | 0.514 | -0.31 | 0.756 |
| Clover<br>(Spring) | total visits |  | Ratio_250_log | -0.655 | 0.083 | -7.92 | <0.001 |
|  | honeybee visits |  | Ratio_250_log | -1.024 | 0.187 | -5.48 | <0.001 |
|  | non-honeybee visits |  | Ratio_250_log | -0.568 | 0.083 | -6.84 | <0.001 |
|  | morphospecies richness |  | Ratio_250_log | -0.290 | 0.053 | -5.46 | <0.001 |
|  | Mean Seed Mass 2022 |  | Ratio_250_log | -0.000 | 0.000 | -2.44 | 0.015 |
|  | Total Seeds 2022 |  | Ratio_250_log | -0.165 | 0.105 | -1.57 | 0.116 |
|  | Mean Mass Individual seedpod 2024 |  | Ratio_250_log | -0.000 | 0.000 | -0.77 | 0.442 |
|  | HTP |  | Ratio_250_log | -0.159 | 0.514 | -0.31 | 0.756 |

Consistent with H1b, the composition of pollinators on clover significantly varied with I:N ratio (df=1, SumofSquares=3.12, R2= 0.0669, F= 9.75, p=0.001) as did pollinator multivariate abundance (residual degrees of freedom = 136, deviance= 281.20, p=0.001). Based on our multivariate abundance analysis, these changes in composition were driven by decreased abundance of honeybees (deviance=13.62, p=0.016) and several species of native medium to long-tongued bees on clover with increasing I:N ratio. These decreasing bees included the yellow-faced bumblebee (*Bombus vosnesenskii*) (deviance=35.56, p=0.001;), the black-tailed bumblebee (*B. melanopygus edwardsii*) (deviance=31.54, p=0.001), the California bumblebee (*B. californicus*) (deviance=18.47, p=0.003), the blue mason bee *Osmia cara* (deviance=15.46, p=0.004;) and an unidentified mason bee (*Osmia sp.*) (deviance=16.95, p=0.003) The common ringlet butterfly (*Coenonympha tullia*) likewise decreased in abundance on clover with increasing I:N ratio (deviance=21.64, p=0.002). All of these species also visited invasive vetch within our dataset.

Likewise consistent with H1b, in summer, pollinator morphospecies richness and visitor abundance on the native sunflower patches decreased with increasing I:N ratio (Figure 2E & D, Table 1), as did the total number of honeybee visits (Table 1) and the total number of non-honeybee pollinator visits to sunflower (Table 1). Pollinator community composition on sunflower varied with I:N ratio (df=1, sum of squares = 1.14, R2= 0.02, F=3.47, p=0.001) as did pollinator multivariate abundance (residual degrees of freedom = 209, deviance= 259.9, p=0.001). Based on our multivariate abundance analysis, these changes in composition were driven by decreased abundances of western honeybees (deviance=35.87, p=0.001), the American lady (*Vanessa virginiensis*) (deviance=17.63, p=0.001), the yellow-shouldered drone fly (*Eristalis stipador*) (deviance= 14.62, p=0.012;), and the beetle *Nemognatha scuttelaris* (deviance=12.92, p=0.038. All of these species also visited the invasive thistle with the exception of *V. virginiensis*.

**Figure 2.**
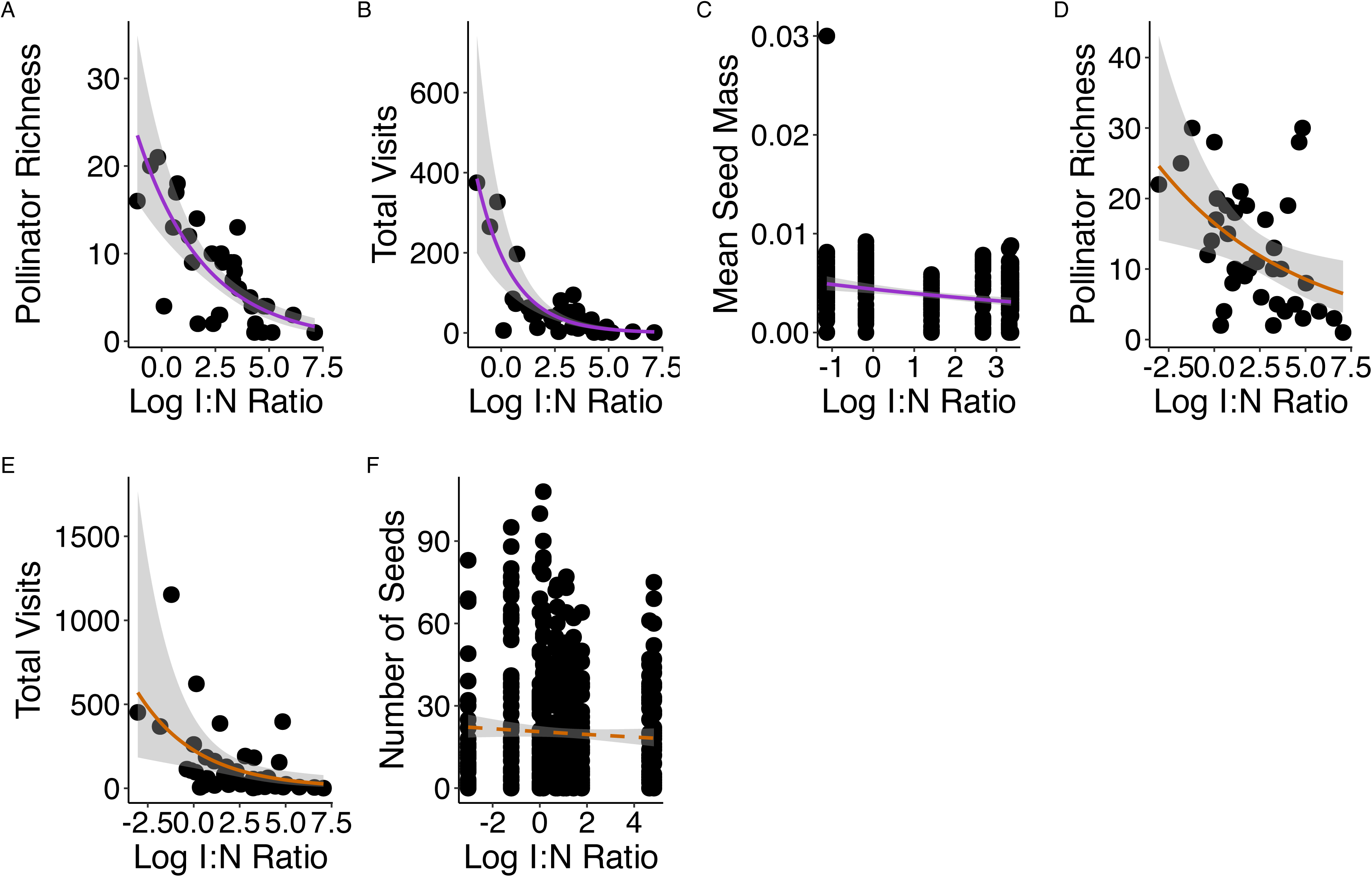
The relationship between the log ratio of invasive plant to native plant floral abundance at 250 M on network metrics for (A) the Müeller indirect effect of clover on vetch, (B) the Müeller indirect effect of vetch on clover, (C) the individual contribution of clover to network nestedness, (D) the individual contribution of vetch to network nestedness, (E) the Müeller indirect effect of sunflower on thistle, (F) the Müeller indirect effect of thistle on sunflower, (G) the individual contribution of sunflower to network nestedness and (H) the individual contribution of thistle to network nestedness. Each dot represents a network for a given meadow in a given season. Solid regression lines denote a statistically significant relationship at an alpha level of 0.05, while dashed regression lines denote a non-significant relationship.

### Visit Quality to Native Plants

Consistent with H1c, heterospecific pollen transfer from non-serpentine invasive plants to native serpentine plants was relatively uncommon. For both native clovers and sunflowers, only 9.72% (7/72) of stigmas counted for both species had invasive plant pollen adhered, and I:N ratio did not correlate with the amount of invasive plant pollen deposited on the stigma (Table 1).

### Native Plant Seed Set

In support of (H2), clover seed set decreased with I:N ratio. The mean seed mass of clover per seedpod also decreased with increasing I:N ratio at 250 m (Table 1; Figure 1C). The total number of clover seeds per pod did not vary with I:N ratio (Table 1). The total number of sunflowers seeds per flowerhead did not significantly change with increasing I:N ratio (Table 1) (Figure 1F).

### Meadow Plant-Pollinator Networks

In support of H3, the functional importance of focal plant species in the network varied with I:N ratio at the meadow scale (Figure 2, Figure S1). In spring, as I:N ratio increased, the indirect effect of clovers on vetch via shared pollinators in the network decreased (Table 2, Figure 2A), but the indirect effect of vetch on clover did not significantly change (Table 2, Figure 2B). Likewise, as the I:N ratio increased, the clover’s individual contribution to network nestedness (Table 2), while those of the vetch did not change (Table 2, Figure 2C). The I:N ratio at the boundary did not significantly correlate with pollinator similarity between vetch and clover based on the Jaccard index of similarity or Sorensen index (Table 2).

**Table 2.**
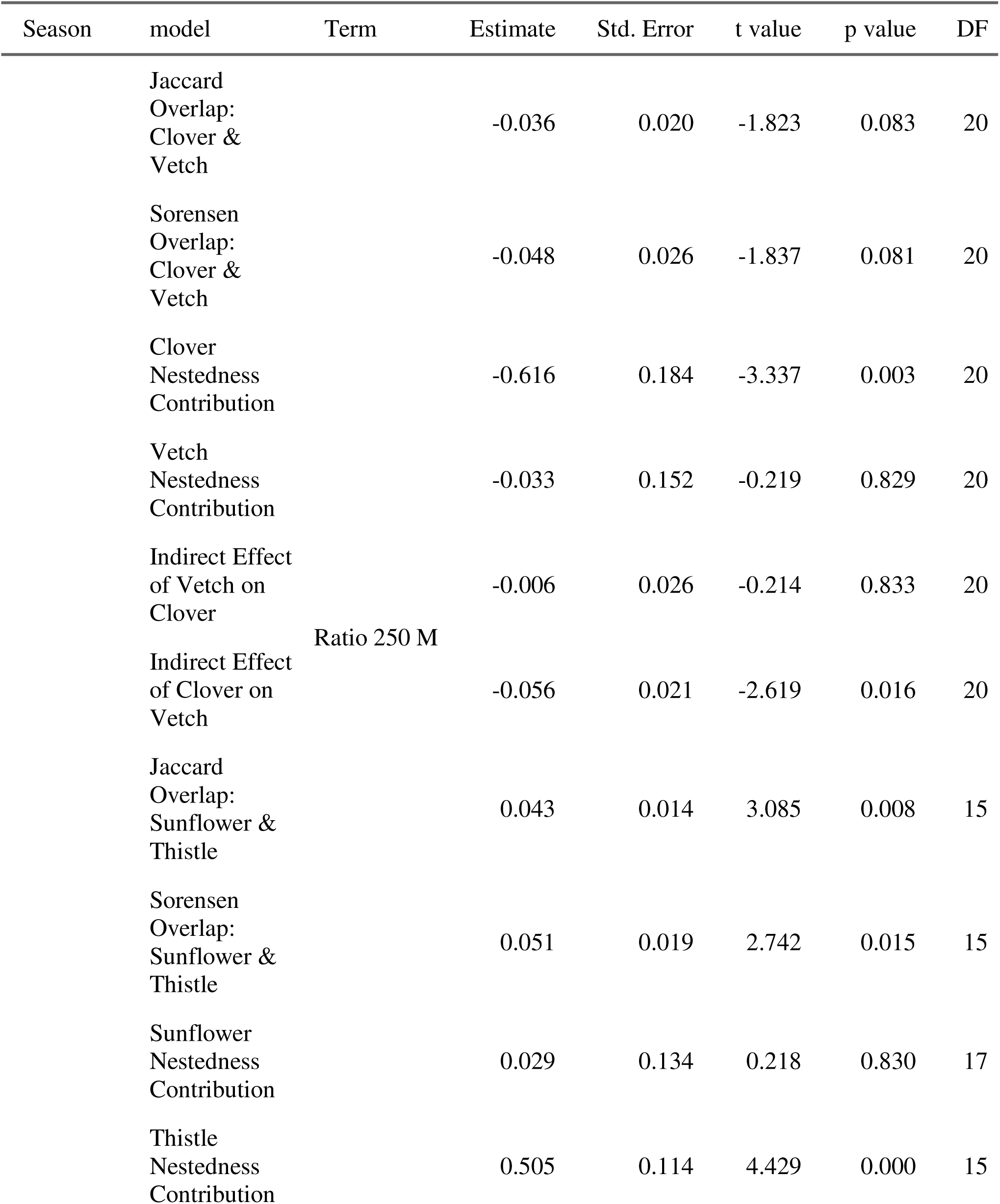
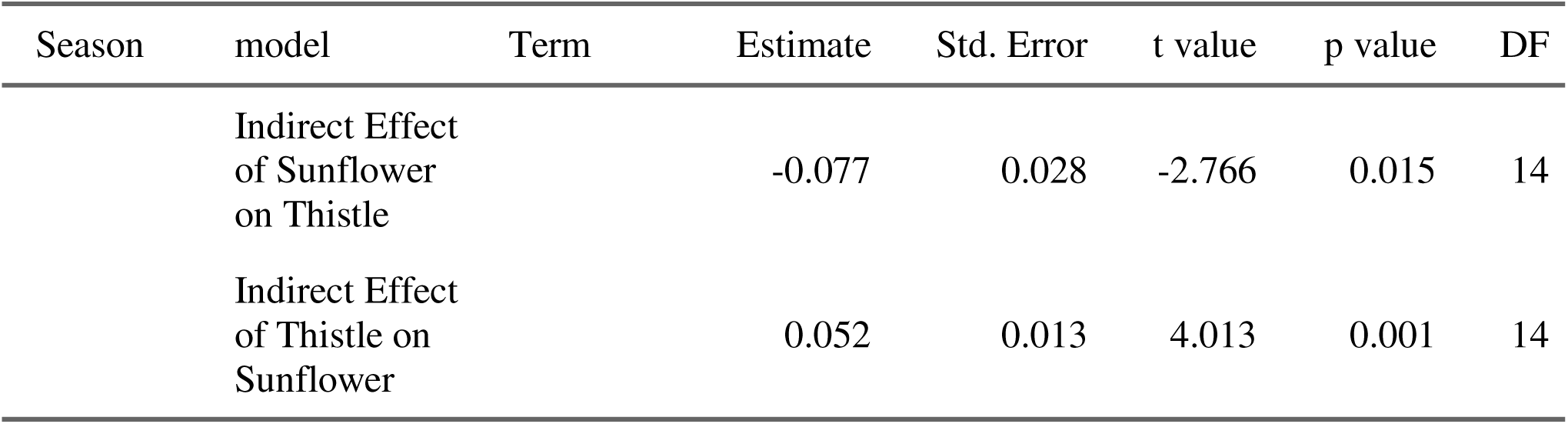
Meadow-scale analyses. Model results for linear models with a fixed effect of invasive to native plant ratio at 250 m on meadow-level properties. Indirect effects were measured using the Müeller index. p<0.05 is the alpha level for statistical significance.

In summer, as the invasive thistle to native sunflower ratio increased, the indirect effect of sunflowers on thistle via shared pollinators in the network as measured by the Müller index decreased (Table 2, Figure 2E), while the invasive thistle’s indirect effects on the sunflower increased (Table 2, Figure 2F). Moreover, as the I:N ratio increased, the individual contribution of sunflower to network nestedness did not change (Table 2, Figure 2G), but the individual contribution of yellow star-thistle to the network’s nestedness increased (Table 2, Figure 2H). As the I:N ratio at the boundary increased, pollinator similarity between thistle and sunflower increased as measured by the Jaccard’s and Sorensen’s indices (Table 2), suggesting increased sharing of pollinators between thistle and sunflower.

## Discussion

As the ratio of invasive non-serpentine flowers to native serpentine flowers at habitat boundaries increased, native plant pollinator visits, pollinator richness, seed set, and native plant’s network role as hubs for pollinators all decreased. These findings support our hypothesis that invasive plants can exert strong indirect effects on native plants across habitat boundaries. Through competition for shared pollinators, invasive plants diminished fitness and disrupted the functional role of native plants in plant-pollinator networks.

Pollinator richness and abundance on focal patches of native plants decreased as the ratio of invasive plants to native plants adjacent to boundaries increased. These results suggest that pollinators are favoring the relatively more abundant plant species when foraging, consistent with theory that pollinators can adaptively track shifts in floral reward availability (Valdovinos et al. 2016, Johnson et al. 2022). Our findings from a wildland system are consistent with prior work showing that the abundance of agricultural crops can affect pollinator spillover into non-crop habitats (Ekroos et al. 2008, Magrach et al. 2017). Although pollinator movement from nesting habitat to foraging habitat explained pollinator spillover at a swamp-prairie boundary (Artz and Waddington 2006), pollinator spillover in our system arose by pollinators foraging for floral resources in adjacent habitats. In contrast to our result, pollinators mostly foraged within their home patches within a naturally fragmented glade habitat (Harmon-Threatt and Anderson 2023). Differences between the two results might be due to borders that are physically more impermeable or at least create greater resistance to movement at glade-wooded borders than in our open grasslands.

Seed set by focal native plants decreased with increasing invasive to native plant ratio at the boundary, suggests that increased spillover competition for shared pollinators across the boundary likely decreases native plant fitness. Within single habitats, invasive plants were shown to have both positive and negative indirect effects on native plant fitness via facilitation of or competition for shared pollinators (Molina-Montenegro et al. 2008, Abdallah et al. 2021, Etter et al. 2022). Whether these indirect effects of shared pollinators are facilitative or competitive may further depend on spatial scale (Ekroos et al. 2008, Underwood et al. 2020). Consistent with prior studies of pollinators spillover in croplands (Tscharntke et al. 2005, Ekroos et al. 2008, Magrach et al. 2017), we found a relationship between the relative plant abundance at the boundary and plant fitness in a wildland system. The mechanisms, however, differed: we found with increasing invasive plant abundance decreased pollinator visitation (including that of honeybees) and seed set, while increasing crop density at a cropland-wildland boundary increased honeybee spillover to native plants, decreasing seed set through decreased visit quality (Magrach et al. 2017).

Although previous studies have found that within a single habitat, invasive plant species can decrease native plant visit quality through increased heterospecific pollen transfer (Arceo-Gómez and Ashman 2016, Parra-Tabla et al. 2021), cross-boundary heterospecific pollen transfer from invasive to native plants was relatively uncommon in our study and did not vary with invasive to native plant ratio at the boundary, perhaps because pollen is typically carried over by pollinators over relatively short distances (e.g. < five meters, Waser and Price 1984, Cresswell et al. 1995, Hung et al. 2023)) and deposited mostly on the first few plants visited (Harder and Barrett 1996, Richards et al. 2009).

As the ratio of invasive to native plants increased at the boundary, the functional importance of native plant species as core hubs for pollinators in the plant-pollinator network decreased, while that of invasive plant species increased. This finding is consistent studies from within single habitats where invasive plant species became highly integrated into the plant-pollinator network, functionally replacing native species as core hubs for pollinators (Russo et al. 2019, Parra-Tabla and Arceo-Gómez 2021). Theory suggests that fewer, but higher quality pollinator visits can allow animal-pollinated species to invade, eventually becoming dominant network hubs (Valdovinos et al. 2009, 2023). Similar to our findings, pollinator spillover across an agricultural boundary altered network structure through decreasing pollinator niche breadth as a result of increased apparent competition between honeybees and other pollinators (Magrach et al. 2017).

Consistent with findings of pollinator spillover in croplands (Ekroos et al. 2008, Tscharntke et al. 2012) and theory (Amarasekare 2004), our findings further suggest that spillover competition for shared pollinators can affect pollinator visitation and plant fitness in wildland systems. While prior work has shown that invasive species can have pervasive cross-boundary effects on food webs and nutrient cycling (Peller and Altermatt 2024), our work demonstrates that invasive plant species can have cross-boundary effects on plant-pollinator mutualisms. Spillover competition may have downstream effects on the biodiversity, function and ecosystem services of plant-pollinator mutualisms (Magrach et al. 2017, Scherer-Lorenzen et al. 2022), raising the salience of accounting for cross-boundary effects when managing invasive species and restoring habitats. Areas thought to be spatial refugia may still be impacted by invasive plants. Examining whether spillover modulates community responses to anthropogenic perturbations can inform the conservation of plant-pollinator mutualisms.

## Acknowledgements

Thank you to Cathy Koehler, Paul Aigner, and Ben Amann at the McLaughlin Reserve for providing logistical support and help with site selection. The research was supported by the contributions of the following individuals as field and lab technicians: Bita Rostami, Alexis Grana, Isabel Mendoza, Nat Walts, Rebekah Shane, Kyle Bianchi, Izzie Wolter, KT Lynch, Seth Tantuico, Ngoc Thu Tat, Monica Rodriguez, Ryan Li, Anastasia Karp, Zach Schneider, and Alison Appelgate. At the Bohart Museum of Entomology, Thomas Zavortink identified bee taxa and Socrates Letana identified fly taxa. Nick Haddad, the UC Davis Ecology Statistics Support Group, Elizabeth Crone and members of the Valdovinos Lab for provided valuable feedback on data analysis. Megan Vahsen provided useful feedback on the manuscript.

## Author Contributions

Study Design & Conceptualization: Susan Harrison, Neal Williams, Rebecca Nelson, Fernanda Valdovinos; Data curation: Rebecca Nelson; Formal analysis: Rebecca Nelson; Methodology: Susan Harrison, Neal Williams, Rebecca Nelson, Fernanda Valdovinos; Visualization: Rebecca Nelson, Neal Williams, Susan Harrison; Writing - original draft: Rebecca Nelson; Writing - review & editing: Rebecca Nelson, Susan Harrison, Neal Williams, Fernanda Valdovinos

