## Supplemental Figures and Tables for "Invasive plants affect native plant pollination through pollinator-mediated cross-boundary effects"

**Supplemental Methods**

We determined the edges of a patch by finding the point at which the nearest conspecific neighboring individual plants were greater than one meter away. To map the patch as a geospatial polygon, we then walked slowly around the edges of the patch, using the CalFlora application to mark the locations of the edges of the patch, generating a polygon in the shape of the patch. We uploaded the geospatial map of plant patch polygons into Google Earth to confirm that the location of the patches was accurate by comparing landscape features in satellite imagery to photographs taken of each meadow in the field. We used ground-truthing to confirm that CalFlora polygons were realistic in their measurements of patch area and extent by measuring the length and width of the patches in the field using a tape measure to estimate the area of patch. If an individual focal plant was greater than one meter from its nearest conspecific neighbor, it was considered an isolated individual and not part of the patch, and therefore plant-pollinator interactions were not observed at it.

We uploaded our plant geospatial data from CalFlora into Google Earth and then used the shape feature to draw 250m, 500m, 1000m radii in relation to the centroid of each focal native plant patch. We combined geospatial information on plant patch locations with patch-level floral abundance data from plant abundance surveys described below to determine the mean abundance of floral units of the invasive plant within fixed 250m, 500m and 1000m radii of a given focal patch of native plants. To determine these radii, we uploaded our plant geospatial data from CalFlora into Google Earth and then used the shape feature to draw 250m, 500m, 1000m radii in relation to the centroid of each focal native plant patch.

To test whether pollinator visitation on focal native plants decreased with raw invasive plant floral abundance, calculated as the mean invasive plant floral abundance within a given spatial radii, we fit generalized linear models with negative binomial distributions for all pollinator response variables and linear models for seed variables using lme4 R package (Bates et al. 2009). We fit separate models for each spatial radii (250 m, 500 m, 1000 m radii) and season (spring and summer) at both the individual flowerhead and patch scales. We further repeated all network and similarity analyzes using generalized linear models with a fixed of invasive plant abundance.

To further account for the effects of variation in native plant patch size on visitation variables, we normalized the ratio of invasive plant floral abundance to native plant floral abundance as follows:

$$Eqn. S1: Area-Normalized Invasive to Native Plant Ratio= \frac{Floral Abundance Invasive Plant}{Floral Abundance Native Plant}*(\frac{Patch Area Native Plant}{\pi\left( {invasive plant radius}^{2} \right).}$$

We took the natural log of area-normalized invasive to native plant ratio and fit negative binomial models with a fixed effect of area-normalized invasive to native plant ratio on visitation response variables (pollinator visitation, honeybee visitation, non-honeybee visitation, pollinator morphospecies richness) for each focal radii and season.

We repeated visitation, heterospecific pollen, and seed variables analyses at the 500 m and 1000 m radii, fitting models with a fixed of natural log-transformed invasive to native plant ratio at 250 m and a random effect of meadow-year. We repeated visitation variables analyses at the individual flowerhead scale, fitting models with a fixed of natural log-transformed invasive to native plant ratio at 250 m and a random effect of meadow-year.

Although the amount of time spent in each meadow was allowed to vary with insect and ﬂoral abundance, each floral unit was sampled for the same amount in order to standardize the sampling effort per ﬂower over a standardized unit of area (Lopezaraiza–Mikel et al. 2007). When measuring visits and seed set to individual flowerheads, we noted cases where focal flowerheads went missing or were eaten by florivores.

**Supplemental Results**

***Invasive Plant Abundance***

Sunflower and clover visitation variables and heterospecific pollen counts did not correlate with natural-log transformed mean invasive plant abundance at 250 m (Table S5). While sunflower seed set did not correlate with invasive plant abundance, clover seed mass in 2024 was positively correlated with invasive plant abundance (Table S5).

The index of Jaccard similarity in pollinators between clover and vetch did not significantly vary with vetch abundance at 250 m (Table S6). Likewise, the Sorensen index of similarity in pollinators between clover and vetch did not significantly vary with vetch abundance at 250 m (Table S6).The index of Jaccard similarity in pollinators between sunflower and thistle significantly increased with thistle abundance at 250 m (Table S6). The index of Sorensen similarity in pollinators between sunflower and thistle increased with increasing thistle abundance at 250 m (Table S6).

At the network scale (250 m), betweenness centrality of clover in the plant-pollinator network was not significantly related to vetch abundance (Table S6), nor was betweenness centrality of vetch (Table S6). Likewise closeness centrality of clover in the network was not significantly related to vetch abundance (Table S6), nor was closeness centrality of vetch (Table S6). The individual contribution of clover to network nestedness did not vary with vetch abundance (Table S6), nor did the individual contribution of vetch to nestedness (Table S6). The indirect effect of vetch on clover via apparent competition for shared pollinators (Müeller Index) significantly increased with vetch abundance (Table S6), but the indirect effect of clover on vetch did not vary with vetch abundance (Table S6).

At the network scale (250 m), betweenness centrality of sunflower in the network did not significantly vary with thistle abundance (Table S6), but betweenness centrality of thistle in the network increased with increasing thistle abundance (Table S6). Sunflower closeness centrality in the network did not vary with thistle abundance (Table S6), nor did thistle closeness centrality (Table S6). The individual contribution of sunflower to network nestedness did not vary with increasing thistle abundance (Table S6), while the nestedness contribution of thistle increased (Table S6). The indirect effect of thistle on sunflower via apparent competition for shared pollinators (Müeller Index) increased with increasing thistle abundance (Table S6), while the indirect effect of sunflower on thistle did not significantly vary with thistle abundance (Table S6).

***Area-Normalized Invasive to Native Plant Ratio***

In spring, as the area-normalized invasive to native plant ratio increased, total pollinator visitation to native clovers significantly decreased at 250 m, but other visitation variables did not change (Table S7). Area-normalized ratio did not correlate with clover seed set and heterospecific pollen transfer variables (Table S7). In summer, pollinator visitation to native sunflowers did not significantly vary with area-normalized invasive to native plant ratio, nor did heterospecific pollen counts and seed set of sunflowers (Table S7).

***500 m and 1000 m Radii***

Our findings were relatively robust and consistent when fitting models with 500 m and 1000 m radii to account for the spatial scales at which bees and other pollinators may forage across the landscape (Tables S8-S9).

**Visitation at Individual Flowerhead Scale**

Pollinator visitation at the individual flowerhead scale was not significantly related to invasive to native plant ratio at 250 m (Table S10).

**Supplemental Figures and Tables**

**Table S1 Meadows.** List of unique meadows and years surveyed and the number of plant species and pollinator morphospecies recorded in the network.

| Season | Year | Site | Number Pollinator Species | Number of Plant Species |
| --- | --- | --- | --- | --- |
| Spring | 2022 | Bertha | 79 | 21 |
| Spring | 2022 | Rock | 28 | 8 |
| Spring | 2022 | Quarry | 65 | 21 |
| Spring | 2022 | Goatgrass | 68 | 33 |
| Spring | 2022 | Long | 69 | 20 |
| Spring | 2022 | Aikawa | 65 | 26 |
| Spring | 2023 | Bertha | 58 | 19 |
| Spring | 2023 | Quarry_1 | 81 | 27 |
| Spring | 2023 | Rock | 20 | 10 |
| Spring | 2024 | Upper_Grid_2 | 18 | 6 |
| Spring | 2024 | Felch | 33 | 8 |
| Spring | 2024 | Coyote | 31 | 11 |
| Spring | 2024 | Aikawa | 36 | 9 |
| Spring | 2024 | Goatgrass | 54 | 15 |
| Spring | 2024 | South Goatgrass | 37 | 6 |
| Spring | 2024 | Anu | 54 | 14 |
| Spring | 2024 | Quarry_1 | 42 | 8 |
| Spring | 2024 | Quarry_2 | 39 | 8 |
| Spring | 2024 | Quarry_3 | 23 | 6 |
| Spring | 2024 | Quarry_4 | 21 | 9 |
| Spring | 2024 | Quarry_5 | 47 | 9 |
| Summer | 2022 | Bertha | 38 | 12 |
| Summer | 2023 | Bertha | 61 | 15 |
| Summer | 2024 | Bertha | 43 | 6 |
| Summer | 2022 | Pond | 40 | 7 |
| Summer | 2023 | Pond | 50 | 13 |
| Summer | 2024 | Pond | 34 | 4 |
| Summer | 2022 | Randy | 29 | 10 |
| Summer | 2023 | Randy | 32 | 9 |
| Summer | 2022 | Aikawa | 38 | 15 |
| Summer | 2022 | Vineyard | 30 | 10 |
| Summer | 2024 | Vineyard | 41 | 8 |
| Summer | 2022 | Lower_Banana | 43 | 10 |
| Summer | 2022 | Banana_Slug | 38 | 11 |
| Summer | 2022 | Quarry Close | 43 | 14 |
| Summer | 2023 | Quarry Close | 39 | 6 |
| Summer | 2024 | Quarry Close | 26 | 7 |
| Summer | 2022 | Quarry Far | 24 | 8 |
| Summer | 2023 | Quarry Far | 42 | 7 |
| Summer | 2024 | Quarry Far | 10 | 2 |

### Table S2. Plant Species List.

| Abbreviation | Plant Species |
| --- | --- |
| AC | Acmispon sp. |
| ACMO | Achyrachaena mollis |
| ACMA | Acmispon americanus |
| ACMI | Achillea millefolium |
| ACBR | Acmispon brachycarpus |
| ACWR | Acmispon wrangelianus |
| AGHE | Agoseris heterophylla |
| ALAM | [Allium amplectens](https://www.inaturalist.org/taxa/60267) |
| Allium | Allium spp. |
| AMME | Amsinckia menziesii |
| ANAR | Anagallis arvensis |
| ANFI | Ancistrocarphus filagineus |
| ASER | Asclepias eriocarpa |
| ASFA | Asclepias fascicularis |
| ASBR | Astragalus breweri |
| ASJE | Astragalus rattanii jepsonii |
| AS | [Astragalus](https://www.inaturalist.org/taxa/49370) |
| BREL | Brodiaea elegans |
| CAEX | Castilleja exserta |
| CA | Calochortus sp. |
| CAPA | Calycadenia pauciflora |
| CAAM | Calochortus amabilis |
| CARU | Castilleja rubicundula |
| CADE | Castilleja densiflora |
| CAMI | Castilleja minor |
| CAME | Calandrinia menziesii |
| CALU | Calochortus luteus |
| CASU | Calochortus superbus |
| CAVE | Calochortus vestae |
| CL | Clarkia gracilis_or_purpurea |
| CLGR | Clarkia gracilis ssp. tracyi |
| CLPU | Clarkia purpurea |
| CAPY | Carduus pycnocephalus |
| CUCA | Cuscuta californica |
| COSP | [Collinsia sparsiflora](https://www.inaturalist.org/taxa/50656) |
| CRHI | Cryptantha hispidula |
| DEHE | Delphinium hesperium |
| DEUL | Delphinium uliginosum |
| DUCY | Dudleya cymosa |
| DEVA | Delphinium variegatum |
| ERNU | Eriogonum nudum |
| DICA | Dichelostemma capitatum |
| ERLA | Eriophyllum lanatum |
| EPDE | Epilobium densiflorum |
| ERGU | Erythranthe guttata |
| ESCA | Eschscholzia californica |
| ERCI | Erodium cicutarium |
| ERBO | Erodium botrys |
| EUSP | [Euphorbia spathulata](https://www.inaturalist.org/taxa/58894) |
| GEMO | Geranium mole |
| GEDI | Geranium dissectum |
| GICA | Gilia capitate |
| GRCA | Grindelia camporum |
| GITR | Gilia tricolor |
| HECU | Heliotropium curassavicum |
| HOCA | Horkelia californica |
| HECO | Hemizonia congesta |
| HOVI | Holocarpha virgata |
| HEAR | Heteromeles arbutifolia |
| HOMA | Hoita macrostachya |
| HIIN | Hirschfeldia incana |
| KECK | Sidalcea keckii |
| LACA | Lasthenia californica |
| LENI | Lepidium nitidum |
| LAMI | Lagophylla minor |
| LATSER | Lactuca serriola |
| LERE | Lewisia rediviva |
| LUAL | Lupinus albifrons |
| LULU | Lupinus luteolus |
| LERA | Lessingia ramulosa |
| LOHO | Lomatium hooverii |
| LASE | Layia septentrionalis |
| LIDI | Linanthus dichotomus |
| LUNA | Lupinus nanus |
| LUBI | Lupinus bicolor |
| LUMI | Lupinus microcarpus |
| LUSU | Lupinus succulentus |
| MEIN | Melilotus indicus |
| MICA | Micropus californicus |
| MIDO | Minuartia douglassi |
| MEPO | Medicago polymorpha |
| WYAN | Wyethia angustifolia |
| PLNO | Plagiobothrys nothofulvus |
| PEKE | Perideridia kelloggii |
| PE | Penstemon spp. |
| PHAQ | Phalaris aquatica |
| PRHE | Primula hendersonii |
| PLER | Plantago erecta |
| RILE | Rigiopappus leptocladus |
| RACA | Ranunculus californicus |
| SAVE | Sairocarpus vexillocalyculatus |
| STAL | Stachys albens |
| SCSI | Scutellaria siphocampyloides |
| SI | Sidalcea sp. |
| SEVU | Senecio vulgaris |
| SIBE | Sisyrinchium bellum |
| HEEX | Helianthus exilis |
| TOVE | Toxicoscordion venenosum |
| TOFR | Toxicoscordion fremontii |
| Toxicoscordion | Toxicoscordion spp. |
| TRAL | Trifolium albopurpureum |
| TRER | Triphysaria eriantha |
| TRBI | Trifolium gracilentum_or_bifidum |
| TRFU | Trifolium fucatum |
| TROB | Trifolium obtusiflorum |
| TRIWI | Trifolium willdenovii |
| TRHI | Trifolium hirtum |
| THCA | Thermopsis californica |
| TRLA-Summer | Trichostema laxum |
| TRLA-Spring | Triteleia laxa |
| UNK_PLANT | Unknown plant |
| VIDO | Viola douglasii |
| VIVI | Vicia villosa |
| CESO | Centaurea solstitialis |
| ZETR | Zeltnera tricantha |
| white-daisy | Leucanthemum vulgare |
| URLI | Uropappus lindleyi |
| VIAM | Vicia americana |
| yellow_lomatium | Lomatium utriculatum |
| CHGL | Chaenactis glabriuscula |
| smooth_catsear | Hypochaeris glabra |
| SOAS | Sonchus asper |
| VICSAT | Vicia sativa |
| LODA | Lomatium dasycarpum |
| PHTA | Phacelia tanacetifolia |
| WYGL | Wyethia glabra |

### Table S3. Pollinator morphospecies list for network analyses. The column “ARTH” lists the morphospecies name identified to the highest taxonomic unit for a given pollinator morphospecies. The column “visit_TRFU” indicates whether a pollinator visited *Trifolium fucatum* (“Yes”), didn’t visit *T. fucatum* (“No”), or did not seasonally overlap with *T. fucatum* (“NA”). The column “visit_HEEX” gives information in the same format for whether a pollinator morphospecies visited *Helianthus exilis*. Season indicates whether or not a pollinator morphospecies was observed in “Spring”, “Summer”, or “Both”.

| **ARTH** | **Description** | **visit_TRFU** | **visit_HEEX** | **Season** |
| --- | --- | --- | --- | --- |
| **Bees (Apoidea)** | | | | |
| **Agapostemon_subtilior** | **Agapostemon_subtilior** | No | Yes | Both |
| **Andrena_baeriae** | **Andrena_baeriae** | Yes | NA | Spring |
| **Andrena_dissona** | **Andrena_dissona** | No | NA | Spring |
| **Andrena_duboisi** | **Andrena_duboisi** | No | NA | Spring |
| **Andrena_nigrocaerulea** | **Andrena_nigrocaerulea** | No | NA | Spring |
| **Andrena_novo_sp.** | Brownish Andrena species undescribed to science. A voucher specimen was submitted to the Bohart Museum of Entomology taxonomists for species description. | NA | No | Summer |
| **Andrena_pallidifovea** | **Andrena_pallidifovea** | Yes | NA | Spring |
| **Andrena_pensilis** | **Andrena_pensilis** | Yes | NA | Spring |
| **Andrena_plana** | **Andrena_plana** | No | NA | Spring |
| **Andrena_sp.** | Unknown striped Andrena species that did not match any of the other species. | No | NA | Spring |
| **Andrena_subchalybea** | **Andrena_subchalybea** | Yes | NA | Spring |
| **Andrena chalybioides** | **Andrena chalybioides** | No | NA | Spring |
| **Anthidium_edwardsii** | **Anthidium_edwardsii** | NA | No | Summer |
| **Anthidium_illustre** | **Anthidium_illustre** | No | Yes | Both |
| **Anthophora_californica** | **Anthophora_californica** | Yes | NA | Spring |
| **Anthophora_crotchii** | **Anthophora_crotchii** | Yes | NA | Spring |
| **Anthophora_edwardsii** | **Anthophora_edwardsii** | No | NA | Spring |
| **Anthophora_sp.** | **Unknown Anthophora_sp. Fuzzy gray *Anthophora* bee.** | NA | No | Summer |
| **Anthophorula_chionura** | **Anthophorula_chionura** | NA | No | Summer |
| **Apis_mellifera** | **Apis_mellifera** | Yes | Yes | Both |
| **Bombus_melanopygus** | **Bombus_melanopygus** | Yes | Yes | Both |
| **Bombus_californicus** | **Bombus_californicus** | Yes | Yes | Both |
| **Bombus_crotchii** | **Bombus_crotchii** | Yes | Yes | Both |
| **Bombus_sp.** | **Unknown Bombus_sp.** | Yes | Yes | Both |
| **Bombus_vosnesenskii** | **Bombus_vosnesenskii** | Yes | Yes | Both |
| **Ashmeadiella_aridula_astragali** | **Ashmeadiella_aridula_astragali** | NA | Yes | Summer |
| **Brachynomada_melanantha** | **Brachynomada_melanantha** | NA | Yes | Summer |
| **Brachysomida_californica** | **Brachysomida_californica** | No | No | Both |
| **Ceratina_punctigena** | **Ceratina_punctigena** | NA | Yes | Summer |
| **Coelixys_editus** | **Coelixys_editus** | NA | Yes | Summer |
| **Diadasia_bituberculata** | **Diadasia_bituberculata** | No | NA | Spring |
| **Diadasia_nigrifrons** | **Diadasia_nigrifrons** | Yes | NA | Spring |
| **Diadasia_sp.** | **Unknown *Diadasia* species, dark gray and fuzzy.** | NA | No | Summer |
| **Dianthidium_dubium** | **Dianthidium_dubium** | NA | Yes | Summer |
| **Duforea sp.** | **Duforea sp.** | No | NA | Spring |
| **Eucera_actuosa** | **Eucera_actuosa** | Yes | NA | Spring |
| **Eucera_frater_albopilosa** | **Eucera_frater_albopilosa** | Yes | No | Both |
| **Eucera_sp.** | **Unknown greenish Eucera_species** | Yes | NA | Spring |
| **Habropoda_depressa** | **Habropoda_depressa** | No | NA | Spring |
| **Habropoda_sp.** | Large brown Habropoda speices that was neither depressa or trissema. | No | NA | Spring |
| **Habropoda_trissema** | **Habropoda_trissema** | Yes | NA | Spring |
| **Halictidae** | **Unknown Halictidae species. Small, metallic striped sweat bee.** | NA | Yes | Summer |
| **Halictus_farinosus** | **Halictus_farinosus** | NA | Yes | Summer |
| **Halictus_ligatus** | **Halictus_ligatus** | No | Yes | Both |
| **Halictus_tripartitus** | **Halictus_tripartitus** | No | Yes | Both |
| **Hoplitus_hypocrita** | **Hoplitus_hypocrita** | No | NA | Spring |
| **Lasiglossum (Hemihalictus sp.)** | **Lasiglossum (Hemihalictus sp.)** | No | NA | Spring |
| **Lasioglossum (Sphecodogastra) sp.** | **Lasioglossum (Sphecodogastra) sp.** | NA | No | Summer |
| **Lasioglossum_helianthi** | **Lasioglossum_helianthi** | NA | No | Summer |
| **Lasioglossum_incompletum** | **Lasioglossum_incompletum** | Yes | Yes | Both |
| **Lasioglossum_titusi** | **Lasioglossum_titusi** | No | Yes | Both |
| **Megachile_apicalis** | **Megachile_apicalis** | NA | Yes | Summer |
| **Megachile_frugalis_pseudofrugalis** | **Megachile_frugalis_pseudofrugalis** | NA | No | Summer |
| **Megachile_montivaga** | **Megachile_montivaga** | NA | Yes | Summer |
| **Megachile_parallela** | **Megachile_parallela** | NA | Yes | Summer |
| **Megachile_sp.** | **Unknown Megachile_species, gray with dark stripes.** | NA | Yes | Summer |
| **Melissodes_lupina** | **Melissodes_lupina** | NA | Yes | Summer |
| **Melissodes_robustior** | **Melissodes_robustior** | NA | Yes | Summer |
| **Nomada_hesperia** | **Nomada_hesperia** | No | NA | Spring |
| **Nomada_obscurella** | **Nomada_obscurella** | No | No | Both |
| **Nomada_vegana** | **Nomada_vegana** | NA | Yes | Summer |
| **Osmia_aglaia** | **Osmia_aglaia** | No | NA | Spring |
| **Osmia_atrocyanea** | **Osmia_atrocyanea** | No | NA | Spring |
| **Osmia_cara** | **Osmia_cara** | Yes | NA | Spring |
| **Osmia_nemoris** | **Osmia_nemoris** | Yes | No | Both |
| **Osmia_sanctaerosae** | **Osmia_sanctaerosae** | No | NA | Spring |
| **Osmia_sp.** | Unknown bluish black metallic Osmia species | Yes | NA | Spring |
| **Panurginus_sp.** | **Panurginus_species** | Yes | No | Both |
| **Peponapis_sp.** | **Peponapis_species** | NA | No | Summer |
| **Triepeolus_utahensis** | **Triepeolus_utahensis** | NA | No | Summer |
| **Xeromelecta_californica** | **Xeromelecta_californica** | NA | Yes | Summer |
| **Xylocopa_sonorina** | **Xylocopa_sonorina** | Yes | NA | Spring |
| **Xylocopa_sp.** | **Unknown Xylocopa_sp. Dark and shiny large carpenter bee.** | Yes | Yes | Both |
| **Xylocopa_tabaniformis** | **Xylocopa_tabaniformis** | Yes | No | Both |
| **Wasps** | | | | |
| **Braconidae** | **Small, black Braconidae wasps** | No | Yes | Both |
| **red_Braconidae** | **Red Braconidae wasps** | No | NA | Spring |
| **Chrysis_angolensis** | **Chrysis_angolensis** | No | NA | Spring |
| **Euodyneras_hidalgo** | **Euodyneras_hidalgo** | NA | Yes | Summer |
| **Ichneumonidae** | **Ichneumonidae** | Yes | NA | Spring |
| **Polistes_aurifer** | **Polistes_aurifer** | Yes | Yes | Both |
| **Polistes_dorsalis** | **Polistes_dorsalis** | NA | Yes | Summer |
| **Prionyx_sp.** | **Black Prionyx_sp.** | NA | Yes | Summer |
| **red_Prionyx_sp.** | **Red Prionyx species** | NA | No | Summer |
| **Pseudomasaris_coquilletti** | **Pseudomasaris_coquilletti** | No | NA | Spring |
| **Sphex_ashmeadi** | **Sphex_ashmeadi** | NA | Yes | Summer |
| **Sphex_ichneumoneus** | **Sphex_ichneumoneus** | NA | Yes | Summer |
| **Sphex_sp.** | **Unkown Sphex_sp., large dark wasp.** | Yes | No | Both |
| Butterflies and Moths (Lepidoptera) | | | | |
| **Adela_eldorada** | **Adela_eldorada** | Yes | NA | Spring |
| **Adela_flammeusella** | **Adela_flammeusella** | Yes | NA | Spring |
| **Adela_sp.** | **Unknown Adela_sp. Black and white fairy moth.** | No | NA | Spring |
| **Adela_trigrapha** | **Adela_trigrapha** | Yes | NA | Spring |
| **Annaphila_decia** | **Annaphila_decia** | No | NA | Spring |
| **Anthocharis_sara** | **Anthocharis_sara** | No | NA | Spring |
| **Argynnis** | **Argynnis sp.** | NA | Yes | Summer |
| **Autographa_californica** | **Autographa_californica** | No | Yes | Both |
| **Burnsius_communis** | **Burnsius_communis** | No | NA | Spring |
| **Chlosyne_palla** | **Chlosyne_palla** | No | NA | Spring |
| **Coenonympha_californica** | **Coenonympha_californica** | Yes | No | Both |
| **Colias_eurytheme** | **Colias_eurytheme** | No | Yes | Both |
| **Danaus_plexippus** | **Danaus_plexippus** | NA | Yes | Summer |
| **Erynnis_propertius** | **Erynnis_propertius** | No | Yes | Both |
| **Erynnis_sp.** | **Unkown Erynnis_species; dark gray and shiny in color.** | No | NA | Spring |
| **Erynnis_tristus** | **Erynnis_tristus** | NA | Yes | Summer |
| **Euphydryas_chalcedona** | **Euphydryas_chalcedona** | No | NA | Spring |
| **Glaucophysche_lygdamus** | **Glaucophysche_lygdamus** | Yes | NA | Spring |
| **Heliolonche_modicella** | **Heliolonche_modicella** | Yes | Yes | Both |
| **Heliopetes_ericetorum** | **Heliopetes_ericetorum** | NA | Yes | Summer |
| **Heliothis_phloxiphaga** | **Heliothis_phloxiphaga** | NA | Yes | Summer |
| **Heliothodes_diminutivus** | **Heliothodes_diminutivus** | Yes | Yes | Both |
| **Hemaris_thetis** | **Hemaris_thetis** | NA | Yes | Summer |
| **Icaricia_acmon** | **Icaricia_acmon** | Yes | Yes | Both |
| **Junonia_grisea** | **Junonia_grisea** | Yes | Yes | Both |
| **Lycaenidae_sp.** | **Lycaenidae_species** | No | NA | Spring |
| **Nymphalis_californica** | **Nymphalis_californica** | No | NA | Spring |
| **Ochlodes_sp.** | **Ochlodes_sp.** | NA | Yes | Summer |
| **Phyciodes_mylitta** | **Phyciodes_mylitta** | Yes | Yes | Both |
| **Prosperpinus_clarkiae** | **Prosperpinus_clarkiae** | Yes | NA | Spring |
| **red_Pterophoridae** | Red **Pterophoridae species** | No | No | Both |
| **Pterophoridae** | **White Pterophoridae species** | NA | Yes | Summer |
| **Tharsalea_xanthoides** | **Tharsalea_xanthoides** | NA | No | Summer |
| **Vanessa_cardui** | **Vanessa_cardui** | No | NA | Spring |
| **Vanessa_virginiensis** | **Vanessa_virginiensis** | NA | Yes | Summer |
| **Zerene_eurydice** | **Zerene_eurydice** | Yes | No | Both |
| **big_blue_butterfly_Lepidoptera (Lepidoptera)** | Unknown species of large, blue butterfly | No | NA | Spring |
| **black_white_moth_Lepidoptera (Arctiinae)** | Unknown black and white tiger moth species (Arctiinae) | No | NA | Spring |
| **blue_red_stripe_moth_Lepidoptera** | Unknown species of blue and red striped moth (Lepidoptera) | No | NA | Spring |
| **silver_moth_Lepidoptera** | Small silver-gray to blue gray shiny moth species (Lepidoptera) | Yes | Yes | Both |
| **small_brown_moth_Lepidoptera** | Small brown moth species (Lepidoptera) | No | NA | Spring |
| **brown_moth_Lepidoptera** | Medium brown moth species (Lepidoptera) | Yes | No | Both |
| **small_gray_black_moth_Lepidoptera** | Small gray-black moth species (Lepidoptera) | Yes | NA | Spring |
| **white_moth_Lepidoptera** | White moth species (Lepidoptera) | Yes | Yes | Both |
| **Flies (Diptera)** | | | | |
| **Asilidae** | **Asilidae** | No | No | Both |
| **BF1_Diptera (Diptera)** | Small black to gray Diptera species | Yes | Yes | Both |
| **BF3_Diptera (Diptera)** | Large black Diptera species | No | NA | Spring |
| **Bombylius_major** | **Bombylius_major** | Yes | Yes | Both |
| **Calliphoridae** | **Calliphoridae** | NA | Yes | Summer |
| **Conophorus_fenestratus** | **Conophorus_fenestratus** | Yes | NA | Spring |
| **Cylindromyia sp.** | **Cylindromyia sp.** | No | NA | Spring |
| **Cyrtopogon_sp.** | **Cyrtopogon_sp.** | Yes | NA | Spring |
| **Drosophilidae** | **Drosophilidae** | NA | Yes | Summer |
| **Eristalis_hirta** | **Eristalis_hirta** | No | Yes | Both |
| **Eristalis_sp.** | **Unknown *Eristalis*_species. Amber in color with dark markings.** | NA | No | Summer |
| **Eristalis_stipador** | **Eristalis_stipador** | NA | Yes | Summer |
| **Eristalis_tenax** | **Eristalis_tenax** | NA | Yes | Summer |
| **Eupeodes fumipennis** | **Eupeodes fumipennis** | Yes | Yes | Both |
| **Muscidae** | **Muscidae** | Yes | No | Both |
| **Andrena-mimic_Syrphid (Syrphidae)** | Andrena-mimicking syrphid fly species. Small yellow syrphid with dark stripes. | No | NA | Spring |
| **APME-mimic_Syrphid (Syrphidae)** | Honeybee-mimicking Syrphid fly species, amber brown with dark stripes. | No | NA | Spring |
| **Dianthidium-mimic_Syprhid (Syrphidae)** | Dianthidium-mimicking syrphid fly species. Yellow medium syrphid with spotted abdomen. | NA | Yes | Summer |
| **Megachile-mimic_Syrphidae (Syrphidae)** | Megachile-mimicking Syrphid fly species. Dark gray syrphid with dark stripes. | NA | No | Summer |
| **Nomada-mimic_Syrphidae (Syrphidae)** | Nomada-mimicking Syrphid fly. Small, elongated smooth yellow syrphid with black stripes. | No | NA | Spring |
| **SAB_black_Syrphid (Syrphidae)** | Black Syrphidae species with stripes on abdomen | No | NA | Spring |
| **orange_Syrphidae (Syrphidae)** | Unknown orange syrphid species | NA | No | Summer |
| **brown_Syrphidae (Syrphidae)** | Unknown brown syrphid species | NA | Yes | Summer |
| **small_narrow_ab_Syrphidae** | Small Syrphidae fly species with narrow abdomen | NA | No | Summer |
| **Syrphidae** | **Unknown Syrphidae species that did not match any of the other Syrphid morphospecies, red to black in color** | No | NA | Spring |
| **Scaeva_affinis** | **Scaeva_affinis** | Yes | NA | Spring |
| **Scathophaga_stercoraria** | **Scathophaga_stercoraria** | No | NA | Spring |
| **Silvius_gigantulus** | **Silvius_gigantulus** | No | NA | Spring |
| **Sphaerophoria_sp.** | **Black to gray Sphaerophoria_sp.** | Yes | Yes | Both |
| **reddish_Sphaerophoria_sp.** | Red **Sphaerophoria species** | Yes | NA | Spring |
| **Tachinidae** | Black-gray, black-red, to red Tachinidae species | No | Yes | Both |
| **Villa_sp.** | **Villa_sp.** | NA | Yes | Summer |
| **Xylocopa-mimic_Diptera** | Xylocopa mimicking fly species. Large dark, round fly. | NA | Yes | Summer |
| **gray_Bombyliidae** | Gray fuzzy Bombyillidae species | No | No | Both |
| **red_Diptera** | Unknown red fly species (Diptera) | No | NA | Spring |
| **stripe_fly_Diptera** | Small unknown striped fly (Diptera) | NA | Yes | Summer |
| **tiny_pale_green_Diptera** | Small, pale green fly species (Diptera) | NA | Yes | Summer |
| **Beetles (Coleoptera)** | | | | |
| **Acmaeodera_sp.** | **Acmaeodera_sp.** | Yes | NA | Spring |
| **Anastrangalia_laetifca** | **Anastrangalia_laetifca** | NA | Yes | Summer |
| **Anthrenus_sp.** | **Anthrenus_sp.** | Yes | Yes | Both |
| **Buprestidae** | **Buprestidae** | NA | Yes | Summer |
| **Cantharidae** | **Cantharidae** | Yes | NA | Spring |
| **Chrysochus_colbaltinus** | **Chrysochus_colbaltinus** | NA | No | Summer |
| **Curculionidae** | **Curculionidae** | No | No | Both |
| **Diabrotica_undecimpunctata** | **Diabrotica_undecimpunctata** | Yes | Yes | Both |
| **Elateridae** | **Elateridae** | Yes | NA | Spring |
| **Epicauta_puncticollis** | Epicauta_puncticollis | NA | Yes | Summer |
| **Hippodamia_convergens** | **Hippodamia_convergens** | Yes | No | Both |
| **Listrus_sp.** | **Listrus_sp. is a medium, dark gray *Listrus* beetle species** | No | Yes | Both |
| **Listrus_sp._1** | **Listrus_sp._1 is a small black *Listrus* beetle species** | Yes | Yes | Both |
| **Listrus_sp._2** | **Listrus_sp._2 is a larger light gray *Listrus* beetle species** | Yes | Yes | Both |
| **Malachius_sp.** | **Malachius_species** | Yes | NA | Spring |
| **Mordellidae** | **Mordellidae** | No | No | Both |
| **Nemognatha_scutellaris** | **Nemognatha_scutellaris** | NA | Yes | Summer |
| **Tetraopes_basalis** | **Tetraopes_basalis** | No | No | Both |
| **Trichodes_ornatus** | **Trichodes_ornatus** | No | No | Both |
| **blue_beetle_Coleoptera** | **Unknown blue beetle species** | No | NA | Spring |
| **orange_beetle_Coleoptera** | **Unknown orange beetle species** | No | NA | Spring |
| **Other Insects** | | | | |
| **Thysanoptera** | **Thysanoptera** | No | Yes | Both |
| **Rhaphidioptera** | **Rhaphidioptera** | No | NA | Spring |
| **Chrysoperla_sp.** | **Chrysoperla_sp.** | Yes | NA | Spring |
| **Hummingbirds** | | | | |
| **Calypte_anna** | **Calypte_anna** | No | Yes | Both |

**Table S4. Patch Information.** For each native plant species, unique site and patch and year combinations are given. If a patch is left blank, it refers to the presence of additional isolated individual native plants at the site that were not part of the central patch.

| Species | Site | Patch | Year |
| --- | --- | --- | --- |
| Helianthus exilis | Aikawa | Aikawa main | 2022 |
| Helianthus exilis | Aikawa | Aikawa small | 2022 |
| Helianthus exilis | Banana Slug | Banana S1 | 2022 |
| Helianthus exilis | Banana Slug | Banana S2 | 2022 |
| Helianthus exilis | Bertha | Bertha B1 | 2024 |
| Helianthus exilis | Bertha | Bertha S1 | 2022 |
| Helianthus exilis | Bertha | Bertha S1 | 2023 |
| Helianthus exilis | Bertha | Bertha S1 | 2024 |
| Helianthus exilis | Bertha | Bertha S2 | 2022 |
| Helianthus exilis | Lower Banana | Lower_Banana S1 | 2022 |
| Helianthus exilis | Pond | Pond P1 | 2024 |
| Helianthus exilis | Pond | Pond S1 | 2022 |
| Helianthus exilis | Pond | Pond S1 | 2023 |
| Helianthus exilis | Quarry Close | Quarry Q1S1_extra | 2022 |
| Helianthus exilis | Quarry Close | Quarry Q3S3_upper | 2022 |
| Helianthus exilis | Quarry Far | Quarry Q6S6 | 2022 |
| Helianthus exilis | Quarry Close | Quarry QS1 | 2022 |
| Helianthus exilis | Quarry Close | Quarry QS1_upper | 2022 |
| Helianthus exilis | Quarry Close | Quarry QS2 | 2022 |
| Helianthus exilis | Quarry Close | Quarry QS2_upper | 2022 |
| Helianthus exilis | Quarry Close | Quarry QS3 | 2022 |
| Helianthus exilis | Quarry Close | Quarry QS5 | 2022 |
| Helianthus exilis | Quarry Close | Quarry_3 Q3 | 2024 |
| Helianthus exilis | Quarry Close | Quarry_3 S3 | 2023 |
| Helianthus exilis | Quarry Close | Quarry_4 Q4 | 2024 |
| Helianthus exilis | Quarry Close | Quarry_4 S4 | 2022 |
| Helianthus exilis | Quarry Close | Quarry_4 S4 | 2023 |
| Helianthus exilis | Quarry Far | Quarry_5 S1 | 2023 |
| Helianthus exilis | Quarry Far | Quarry_5 S6 | 2023 |
| Helianthus exilis | Quarry Far | Quarry_7 Q7 | 2024 |
| Helianthus exilis | Quarry Far | Quarry_7 S7 | 2023 |
| Helianthus exilis | Randy | Randy S1 | 2022 |
| Helianthus exilis | Randy | Randy S1 | 2023 |
| Helianthus exilis | Randy | Randy S2 | 2022 |
| Helianthus exilis | Randy | Randy S3 | 2022 |
| Helianthus exilis | Randy | Randy S4 | 2022 |
| Helianthus exilis | Randy | Randy burnt_log | 2022 |
| Helianthus exilis | Randy | Randy extra_patch | 2022 |
| Helianthus exilis | Vineyard | Vineyard S1 | 2022 |
| Helianthus exilis | Vineyard | Vineyard S2 | 2022 |
| Helianthus exilis | Vineyard | Vineyard S3 | 2022 |
| Helianthus exilis | Vineyard | Vineyard S4 | 2022 |
| Helianthus exilis | Vineyard | Vineyard S5 | 2022 |
| Helianthus exilis | Vineyard | Vineyard V1 | 2024 |
| Trifolium fucatum | Aikawa | AI | 2024 |
| Trifolium fucatum | Aikawa | ATR1 | 2022 |
| Trifolium fucatum | Aikawa | ATR2 | 2022 |
| Trifolium fucatum | Anu | A1 | 2024 |
| Trifolium fucatum | Anu_1 | A1 | 2024 |
| Trifolium fucatum | Anu_2 | A2 | 2024 |
| Trifolium fucatum | Anu_3 | A3 | 2024 |
| Trifolium fucatum | Anu_4 | A4 | 2024 |
| Trifolium fucatum | Bertha |  | 2023 |
| Trifolium fucatum | Bertha | BTR1 | 2022 |
| Trifolium fucatum | Bertha | BTR2 | 2022 |
| Trifolium fucatum | Bertha | TRFU2 | 2023 |
| Trifolium fucatum | Coyote | C1 | 2024 |
| Trifolium fucatum | Felch | F1 | 2024 |
| Trifolium fucatum | Goatgrass |  | 2023 |
| Trifolium fucatum | Goatgrass | GG1 | 2024 |
| Trifolium fucatum | Goatgrass | GTR1 | 2022 |
| Trifolium fucatum | Goatgrass | GTR2 | 2022 |
| Trifolium fucatum | Goatgrass | misc | 2022 |
| Trifolium fucatum | Long | LTR1 | 2022 |
| Trifolium fucatum | Long | LTR2 | 2022 |
| Trifolium fucatum | Lower_Grid |  | 2023 |
| Trifolium fucatum | Lower_Grid | TRFU1 | 2023 |
| Trifolium fucatum | Lower_Grid_1 |  | 2023 |
| Trifolium fucatum | Oak | OTR1 | 2022 |
| Trifolium fucatum | Pond |  | 2023 |
| Trifolium fucatum | Quarry | Q3 | 2024 |
| Trifolium fucatum | Quarry | Q4 | 2024 |
| Trifolium fucatum | Quarry | Q5 | 2024 |
| Trifolium fucatum | Quarry | QTR1 | 2022 |
| Trifolium fucatum | Quarry | QTR3 | 2022 |
| Trifolium fucatum | Quarry_1 |  | 2023 |
| Trifolium fucatum | Quarry_1 | Q1 | 2024 |
| Trifolium fucatum | Quarry_2 |  | 2023 |
| Trifolium fucatum | Quarry_2 | Q2 | 2024 |
| Trifolium fucatum | Quarry_3 | Q3 | 2024 |
| Trifolium fucatum | Quarry_4 | Q4 | 2024 |
| Trifolium fucatum | Quarry_5 | Q5 | 2024 |
| Trifolium fucatum | Rock |  | 2023 |
| Trifolium fucatum | Rock | R1 | 2024 |
| Trifolium fucatum | Rock | RT1 | 2022 |
| Trifolium fucatum | Rock | TRFU1 | 2023 |
| Trifolium fucatum | Rock | TRFU2 | 2023 |
| Trifolium fucatum | South_Goatgrass | SG1 | 2024 |
| Trifolium fucatum | Tac_Shack | TacTR1 | 2022 |
| Trifolium fucatum | Upper_Grid_2 |  | 2023 |
| Trifolium fucatum | Upper_Grid_2 | TRFU1 | 2023 |
| Trifolium fucatum | Upper_Grid_2 | TRFU2 | 2023 |
| Trifolium fucatum | Upper_Grid_2 | UG2_1 | 2024 |
| Trifolium fucatum | Upper_Grid_2 | UG2_2 | 2024 |

**Table S5.** The statistical outputs of negative binomial and Gaussian for clover seed set mixed effects models, testing for an effect on natural log-transformed 250 m mean invasive plant floral abundance on patch-level visitation, seed set, and heterospecific counts for both spring native bull clovers (*T. fucatum*) and summer native serpentine sunflowers (*H. exilis*). All models had a random of effect of unique meadow-year. Because we used the glmmTMB package, all models had a test statistic of z and thus infinite degrees of freedom associated with the z-distribution. The alpha level for significance was 0.05.

| Species  (Season) | Response | DF | Term | Estimate | Std. Error | z | p-value |
| --- | --- | --- | --- | --- | --- | --- | --- |
| Sunflower  (Summer) | total visits | Infinite | Avg_250_log | -0.107 | 0.146 | -0.73 | 0.463 |
|  | honeybee visits |  |  | -0.197 | 0.214 | -0.92 | 0.356 |
|  | non-honeybee visits |  |  | -0.073 | 0.136 | -0.54 | 0.589 |
|  | morphospecies richness |  |  | -0.006 | 0.070 | -0.09 | 0.932 |
|  | Number of Seeds per seedhead |  |  | 0.030 | 0.059 | 0.51 | 0.611 |
|  | HTP count |  |  | 0.798 | 0.652 | 1.22 | 0.221 |
| Clover  (Spring) | total visits |  |  | -0.131 | 0.236 | -0.55 | 0.580 |
|  | honeybee visits |  |  | 0.184 | 0.360 | 0.51 | 0.609 |
|  | non-honeybee visits |  |  | -0.206 | 0.230 | -0.90 | 0.370 |
|  | morphospecies richness |  |  | -0.085 | 0.112 | -0.76 | 0.445 |
|  | Mean Seed Mass 2022 |  |  | 0.000 | 0.001 | 0.35 | 0.728 |
|  | Total Seeds 2022 |  |  | 0.239 | 0.264 | 0.91 | 0.365 |
|  | Mean Mass Individual seedpod 2024 |  |  | 0.001 | 0.000 | 2.18 | 0.030 |
|  | HTP |  |  | 0.798 | 0.652 | 1.22 | 0.221 |

**Table S6.** Meadow-scale analyses. Model results for linear models with a fixed effect of natural log-transformed mean invasive plant floral abundance at 250 m on meadow-level properties. Indirect effects were measured using the Müeller index. p<0.05 is the alpha level for statistical significance.

| Season | model | Term | Estimate | Std. Error | t value | p value | DF |
| --- | --- | --- | --- | --- | --- | --- | --- |
| Spring | Clover Betweenness Centrality | Avg_250_log | 19.546 | 70.344 | 0.278 | 0.784 | 20 |
|  | Vetch Betweenness Centrality |  | 17.735 | 36.166 | 0.490 | 0.629 | 20 |
|  | Clover Closeness Centrality |  | 0.001 | 0.002 | 0.444 | 0.662 | 20 |
|  | Vetch Closeness Centrality |  | 0.001 | 0.002 | 0.875 | 0.392 | 20 |
|  | Jaccard Overlap: Clover & Vetch |  | 0.054 | 0.037 | 1.458 | 0.160 | 20 |
|  | Sorensen Overlap: Clover & Vetch |  | 0.079 | 0.049 | 1.610 | 0.123 | 20 |
|  | Clover Nestedness Contribution |  | 0.084 | 0.419 | 0.199 | 0.844 | 20 |
|  | Vetch Nestedness Contribution |  | 0.381 | 0.263 | 1.446 | 0.164 | 20 |
|  | Indirect Effect of Vetch on Clover |  | 0.103 | 0.042 | 2.437 | 0.024 | 20 |
|  | Indirect Effect of Clover on Vetch |  | 0.032 | 0.045 | 0.716 | 0.482 | 20 |
| Summmer | Sunflower Betweenness Centrality |  | 30.644 | 31.159 | 0.983 | 0.339 | 17 |
|  | Thistle Betweenness Centrality |  | 57.022 | 14.058 | 4.056 | 0.001 | 15 |
|  | Sunflower Closeness Centrality |  | -0.001 | 0.001 | -0.827 | 0.420 | 17 |
|  | Thistle Closeness Centrality |  | -0.001 | 0.001 | -0.498 | 0.626 | 15 |
|  | Jaccard Overlap: Sunflower & Thistle |  | 0.049 | 0.017 | 2.949 | 0.010 | 15 |
|  | Sorensen Overlap: Sunflower & Thistle |  | 0.059 | 0.022 | 2.710 | 0.016 | 15 |
|  | Sunflower Nestedness Contribution |  | 0.139 | 0.162 | 0.861 | 0.401 | 17 |
|  | Thistle Nestedness Contribution |  | 0.576 | 0.137 | 4.207 | 0.001 | 15 |
|  | Indirect Effect of Sunflower on Thistle |  | -0.066 | 0.037 | -1.776 | 0.098 | 14 |
|  | Indirect Effect of Thistle on Sunflower |  | 0.056 | 0.017 | 3.270 | 0.006 | 14 |

**Table S7.** The statistical outputs of negative binomial and Gaussian for clover seed set mixed effects models, testing for an effect on natural log-transformed invasive to native plant ratio at 250 m normalized by patch area on patch-level visitation, seed set, and heterospecific counts for both spring native bull clovers (*T. fucatum*) and summer native serpentine sunflowers (*H. exilis*). All models had a random of effect of unique meadow-year. Because we used the glmmTMB package, all models had a test statistic of z and thus infinite degrees of freedom associated with the z-distribution. The alpha level for significance was 0.05.

| Species  (Season) | Response | DF | Term | Estimate | Std. Error | z value | p-value |
| --- | --- | --- | --- | --- | --- | --- | --- |
| Sunflower  (Summer) | total visits | Infinite | Ratio_250_area_log | -0.098 | 0.139 | -0.71 | 0.481 |
|  | honeybee visits |  | Ratio_250_area_log | -0.182 | 0.221 | -0.82 | 0.412 |
|  | non-honeybee visits |  | Ratio_250_area_log | -0.065 | 0.109 | -0.60 | 0.549 |
|  | morphospecies richness |  | Ratio_250_area_log | 0.006 | 0.066 | 0.09 | 0.926 |
|  | Number of Seeds per seedhead |  | Ratio_250_area_log | -0.031 | 0.067 | -0.46 | 0.645 |
|  | HTP count |  | Ratio_250_area_log | -0.053 | 0.525 | -0.10 | 0.920 |
| Clover  (Spring) | total visits |  | Ratio_250_area_log | -0.415 | 0.108 | -3.84 | <0.001 |
|  | honeybee visits |  | Ratio_250_area_log | -0.172 | 0.304 | -0.57 | 0.571 |
|  | non-honeybee visits |  | Ratio_250_area_log | -0.179 | 0.118 | -1.51 | 0.131 |
|  | morphospecies richness |  | Ratio_250_area_log | -0.110 | 0.073 | -1.52 | 0.128 |
|  | Mean Seed Mass 2022 |  | Ratio_250_area_log | -0.000 | 0.000 | -1.80 | 0.072 |
|  | Total Seeds 2022 |  | Ratio_250_area_log | -0.157 | 0.114 | -1.38 | 0.169 |
|  | Mean Mass Individual seedpod 2024 |  | Ratio_250_area_log | 0.000 | 0.000 | 1.26 | 0.209 |
|  | HTP |  | Ratio_250_area_log | -0.053 | 0.525 | -0.10 | 0.920 |

**Table S8.** The statistical outputs of negative binomial and Gaussian for clover seed set mixed effects models, testing for an effect on natural log-transformed 500 m invasive to native plant ratio on patch-level visitation, seed set, and heterospecific counts for both spring native bull clovers (*T. fucatum*) and summer native serpentine sunflowers (*H. exilis*). All models had a random of effect of unique meadow-year. Because we used the glmmTMB package, all models had a test statistic of z and thus infinite degrees of freedom associated with the z-distribution. The alpha level for significance was 0.05.

| Species  (Season) | Response | DF | Term | Estimate | Std. Error | z value | p-value |
| --- | --- | --- | --- | --- | --- | --- | --- |
| Sunflower  (Summer) | total visits | Infinite | Ratio_500_log | -0.395 | 0.108 | -3.64 | <0.001 |
|  | honeybee visits |  |  | -0.561 | 0.179 | -3.14 | 0.002 |
|  | non-honeybee visits |  |  | -0.357 | 0.104 | -3.43 | <0.001 |
|  | morphospecies richness |  |  | -0.171 | 0.061 | -2.78 | 0.005 |
|  | Number of Seeds per seedhead |  |  | -0.031 | 0.051 | -0.62 | 0.537 |
|  | HTP count |  |  | -0.233 | 0.386 | -0.60 | 0.545 |
| Clover  (Spring) | total visits |  |  | -0.614 | 0.091 | -6.76 | <0.001 |
|  | honeybee visits |  |  | -0.981 | 0.171 | -5.73 | <0.001 |
|  | non-honeybee visits |  |  | -0.540 | 0.085 | -6.34 | <0.001 |
|  | morphospecies richness |  |  | -0.270 | 0.051 | -5.29 | <0.001 |
|  | Mean Seed Mass 2022 |  |  | -0.000 | 0.000 | -1.51 | 0.132 |
|  | Total Seeds 2022 |  |  | -0.115 | 0.091 | -1.26 | 0.208 |
|  | Mean Mass Individual seedpod 2024 |  |  | -0.000 | 0.000 | -0.04 | 0.968 |
|  | HTP |  |  | -0.233 | 0.386 | -0.60 | 0.545 |

**Table S9.** The statistical outputs of negative binomial and Gaussian for clover seed set mixed effects models, testing for an effect on natural log-transformed 1000 m invasive to native plant ratio on patch-level visitation, seed set, and heterospecific counts for both spring native bull clovers (*T. fucatum*) and summer native serpentine sunflowers (*H. exilis*). All models had a random of effect of unique meadow-year. Because we used the glmmTMB package, all models had a test statistic of z and thus infinite degrees of freedom associated with the z-distribution. The alpha level for significance was 0.05.

| Species  (Season) | Response | DF | Term | Estimate | Std. Error | z value | p-value |
| --- | --- | --- | --- | --- | --- | --- | --- |
| Sunflower  (Summer) | total visits | Infinite | Ratio_1000_log | -0.525 | 0.139 | -3.76 | <0.001 |
|  | honeybee visits |  |  | -0.798 | 0.257 | -3.10 | 0.002 |
|  | non-honeybee visits |  |  | -0.471 | 0.136 | -3.46 | <0.001 |
|  | morphospecies richness |  |  | -0.258 | 0.074 | -3.48 | <0.001 |
|  | Number of Seeds per seedhead |  |  | -0.075 | 0.093 | -0.80 | 0.423 |
|  | HTP count |  |  | -0.271 | 0.329 | -0.82 | 0.410 |
| Clover  (Spring) | total visits |  |  | -0.536 | 0.097 | -5.54 | <0.001 |
|  | honeybee visits |  |  | -0.969 | 0.159 | -6.09 | <0.001 |
|  | non-honeybee visits |  |  | -0.476 | 0.090 | -5.29 | <0.001 |
|  | morphospecies richness |  |  | -0.247 | 0.041 | -5.98 | <0.001 |
|  | Mean Seed Mass 2022 |  |  | -0.000 | 0.000 | -1.76 | 0.078 |
|  | Total Seeds 2022 |  |  | -0.146 | 0.077 | -1.90 | 0.058 |
|  | Mean Mass Individual seedpod 2024 |  |  | 0.000 | 0.000 | 0.11 | 0.916 |
|  | HTP |  |  | -0.271 | 0.329 | -0.82 | 0.410 |

**Table S10.** Individual flowerhead visitation outputs for both spring native bull clovers (*T. fucatum*) and summer native serpentine sunflowers (*H. exilis*). All models had a fixed effect of natural log-transformed invasive to native plant ratio at 250 m and a random of effect of unique meadow-year. Because we used the glmmTMB package, all models had a test statistic of z and thus infinite degrees of freedom associated with the z-distribution. The alpha level for significance was 0.05.

| Species (Season) | Response | Term | Estimate | Std. Error | z value | p-value |
| --- | --- | --- | --- | --- | --- | --- |
| Sunflower  (Summer) | total visits | Ratio_250_log | 0.042 | 0.042 | 0.99 | 0.321 |
|  | honeybee visits |  | 0.021 | 0.092 | 0.23 | 0.819 |
|  | non-honeybee visits |  | 0.046 | 0.040 | 1.15 | 0.249 |
|  | morphospecies richness |  | 0.030 | 0.033 | 0.91 | 0.364 |
| Clover  (Spring) | total visits |  | -0.037 | 0.064 | -0.59 | 0.558 |
|  | honeybee visits |  | -0.940 | 1929842.409 | -0.00 | 1.000 |
|  | non-honeybee visits |  | -0.037 | 0.064 | -0.59 | 0.558 |
|  | morphospecies richness |  | -0.027 | 0.069 | -0.38 | 0.701 |

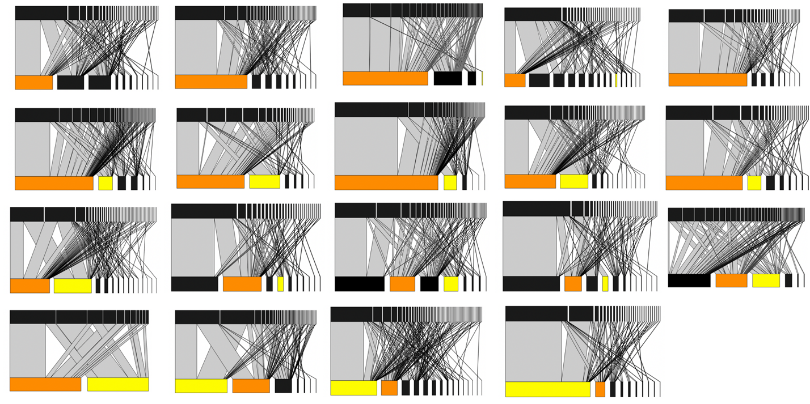

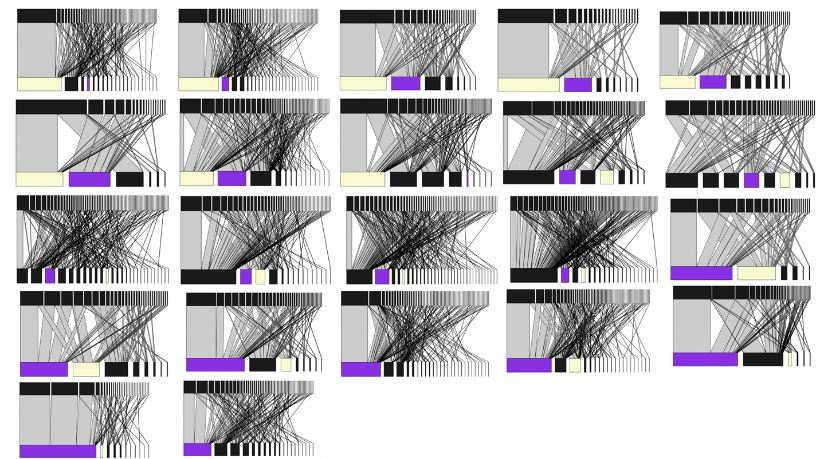
**Figure S1.** Plant-pollinator networks. Each network was constructed for a given meadow and year. The top panel of networks is for spring, and the bottom panel is for summer. The top rows are pollinators, and the bottom rows are plants. Each box represents a different species with the lines between them representing interactions. The thickness of each box and line is proportional the frequency with a species or interaction was observed in the network. For the spring networks, vetch is highlighted in purple and clover in beige. For the summer networks, star-thistle is highlighted in yellow and serpentine sunflower in orange.

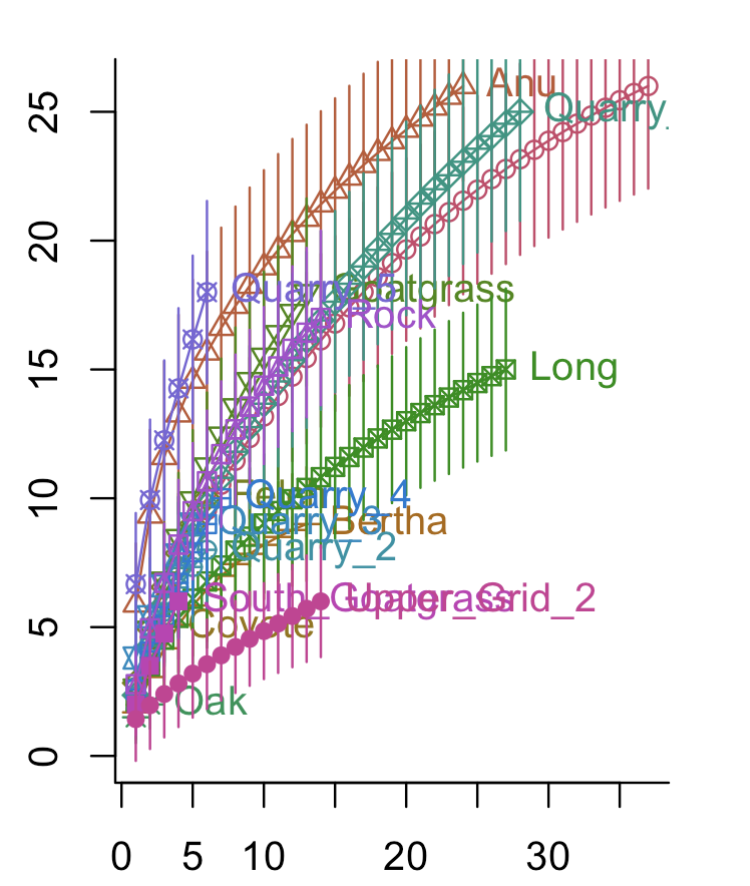

**Figure S2.** Species accumulation curves by meadow for pollinator visitation to *Trifolium fucatum*. The x-axis shows sampling effort, and the y-axis shows pollinator morphospecies richness. Species accumulation curves are colored and labeled by meadow.

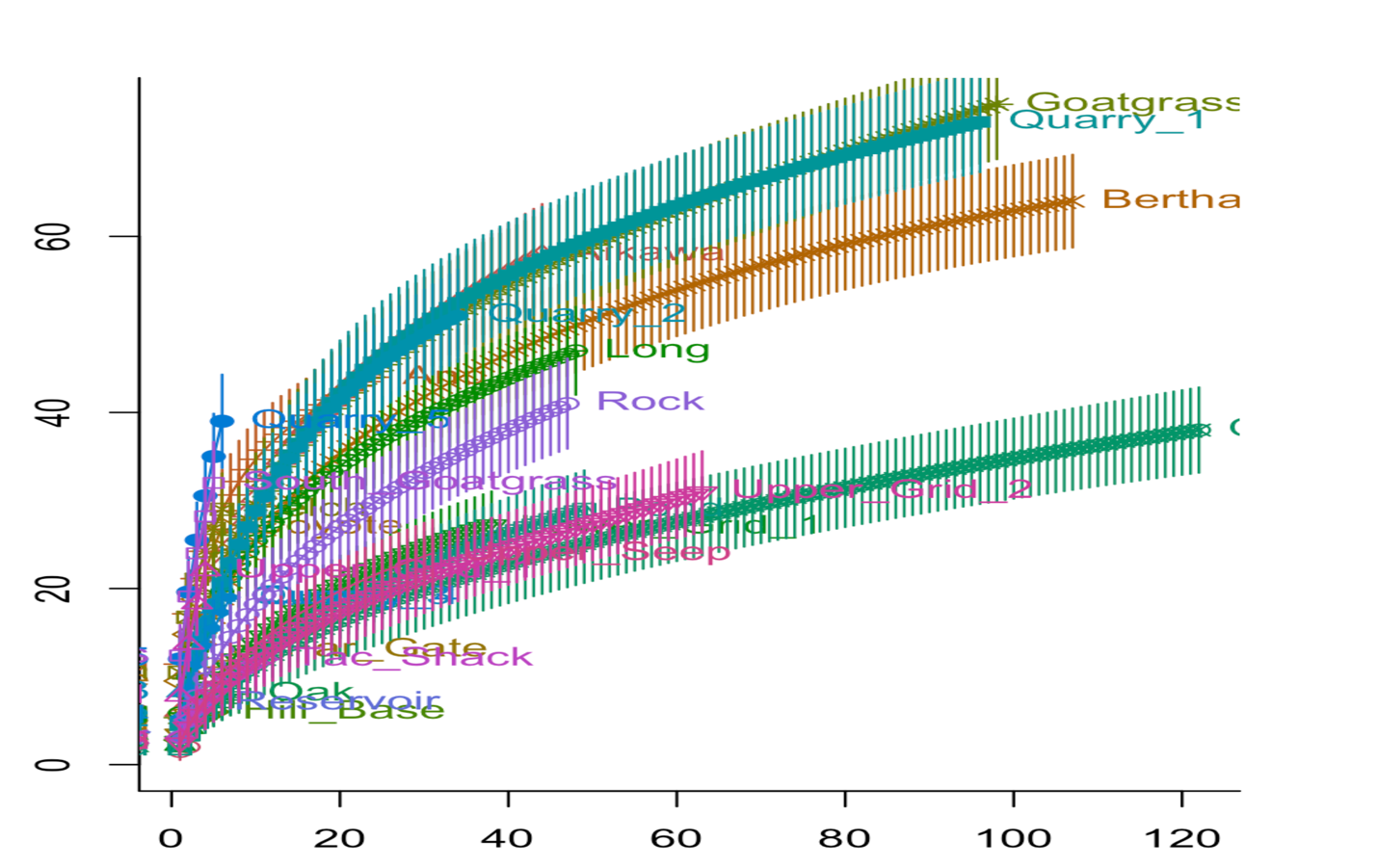

**Figure S3.** Species accumulation curves by meadow for pollinator visitation to full network of spring plants. The x-axis shows sampling effort, and the y-axis shows pollinator morphospecies richness. Species accumulation curves are colored and labeled by meadow.

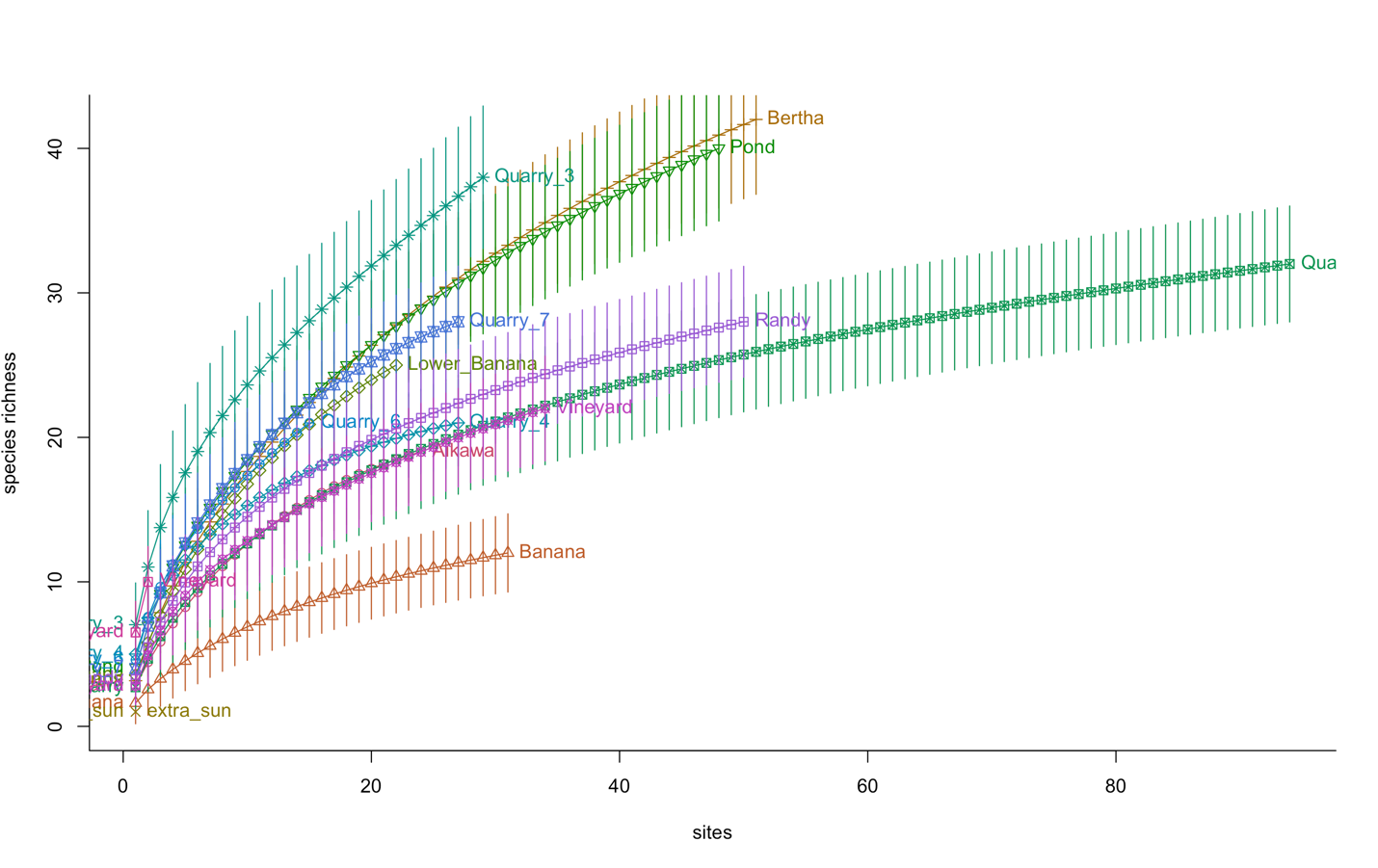

**Figure S4.** Species accumulation curves by meadow for pollinator visitation to *Helianthus exilis*. The x-axis shows sampling effort, and the y-axis shows pollinator morphospecies richness. Species accumulation curves are colored and labeled by meadow.

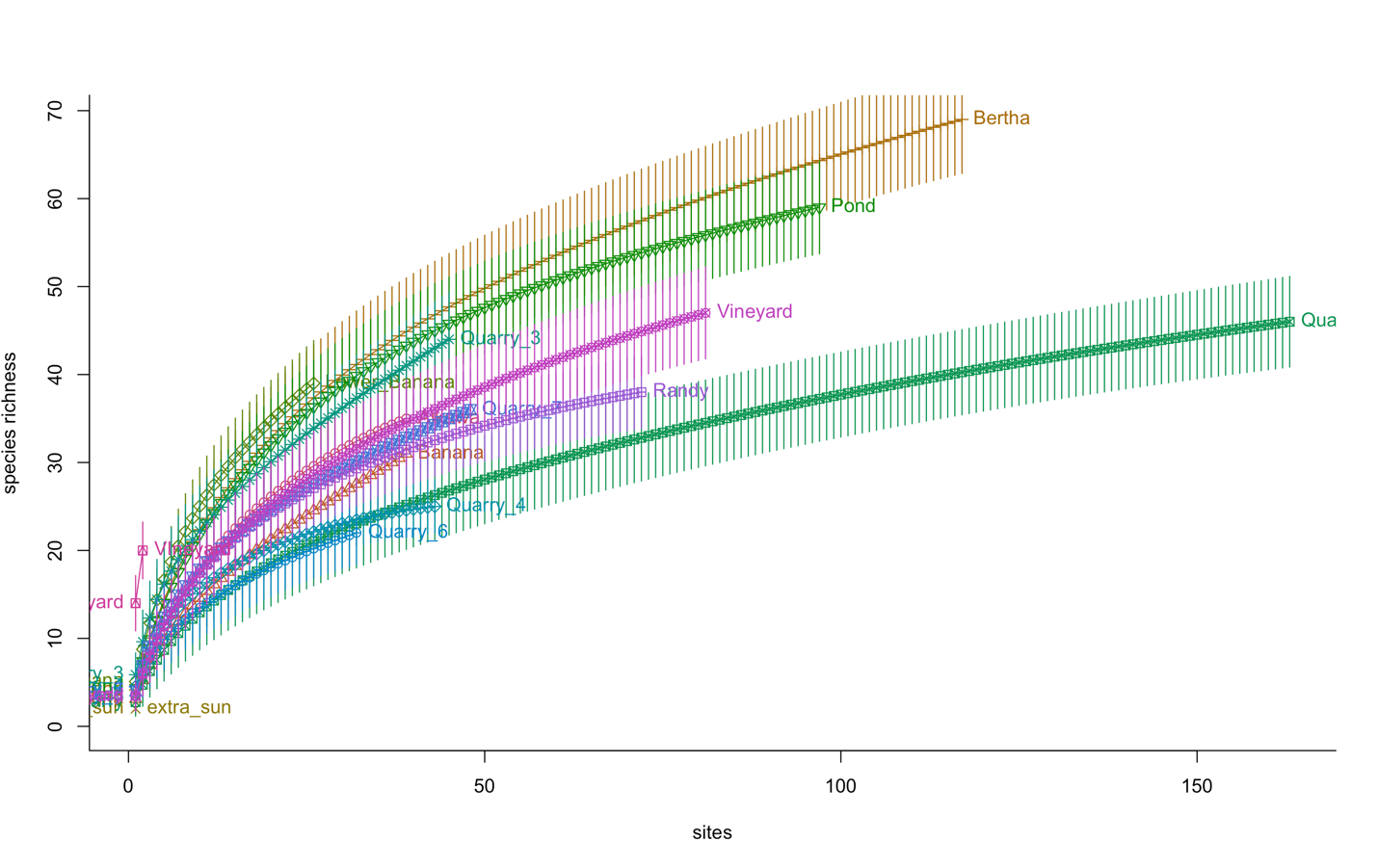

**Figure S5.** Species accumulation curves by meadow for pollinator visitation to full network of summer plants. The x-axis shows sampling effort, and the y-axis shows pollinator morphospecies richness. Species accumulation curves are colored and labeled by meadow.

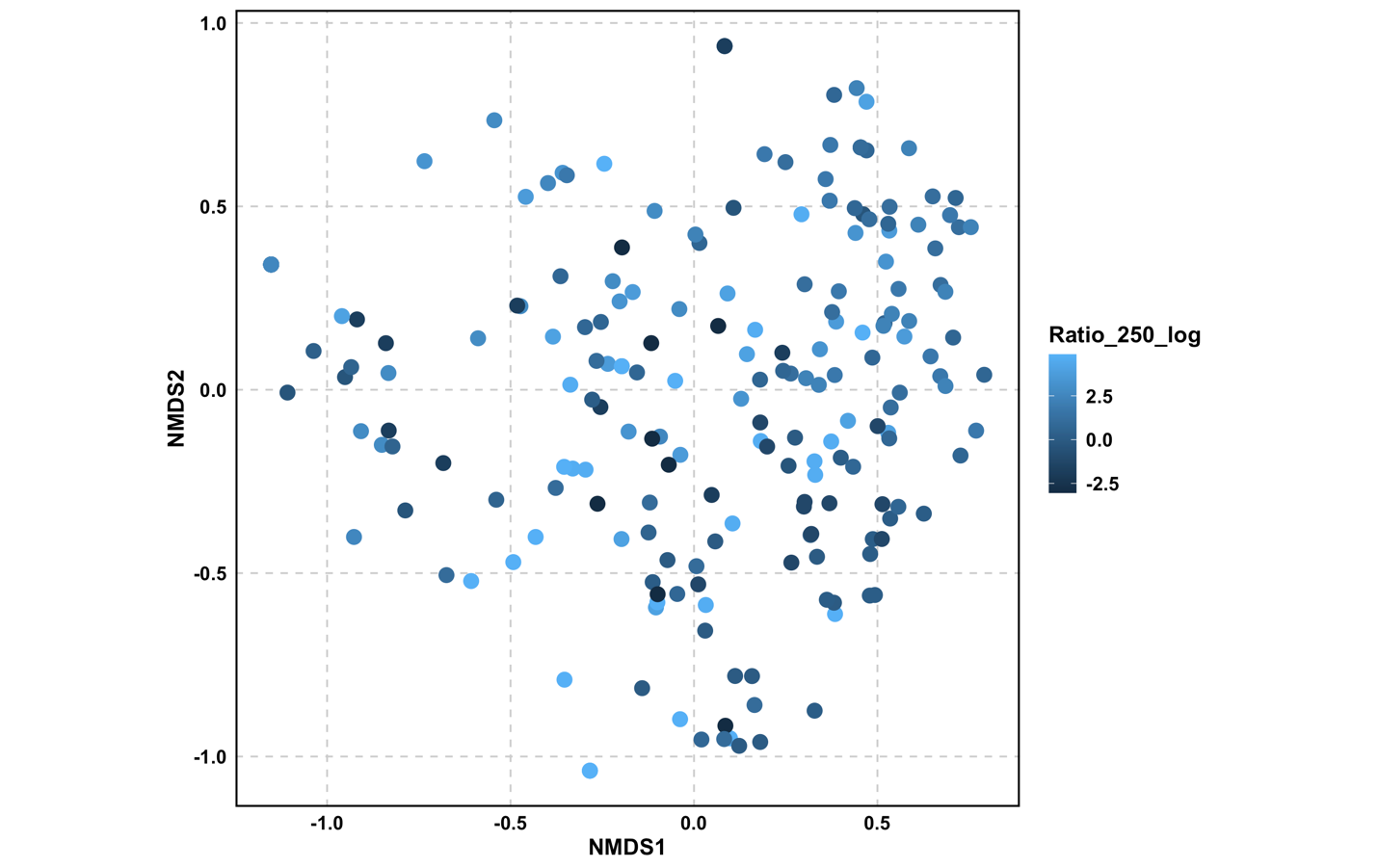

**Figure S6.** NMDS visualization of PERMANOVA of pollinator visitation to *Helianthus exilis* by the log of the invasive to native plant ratio at 250 m.

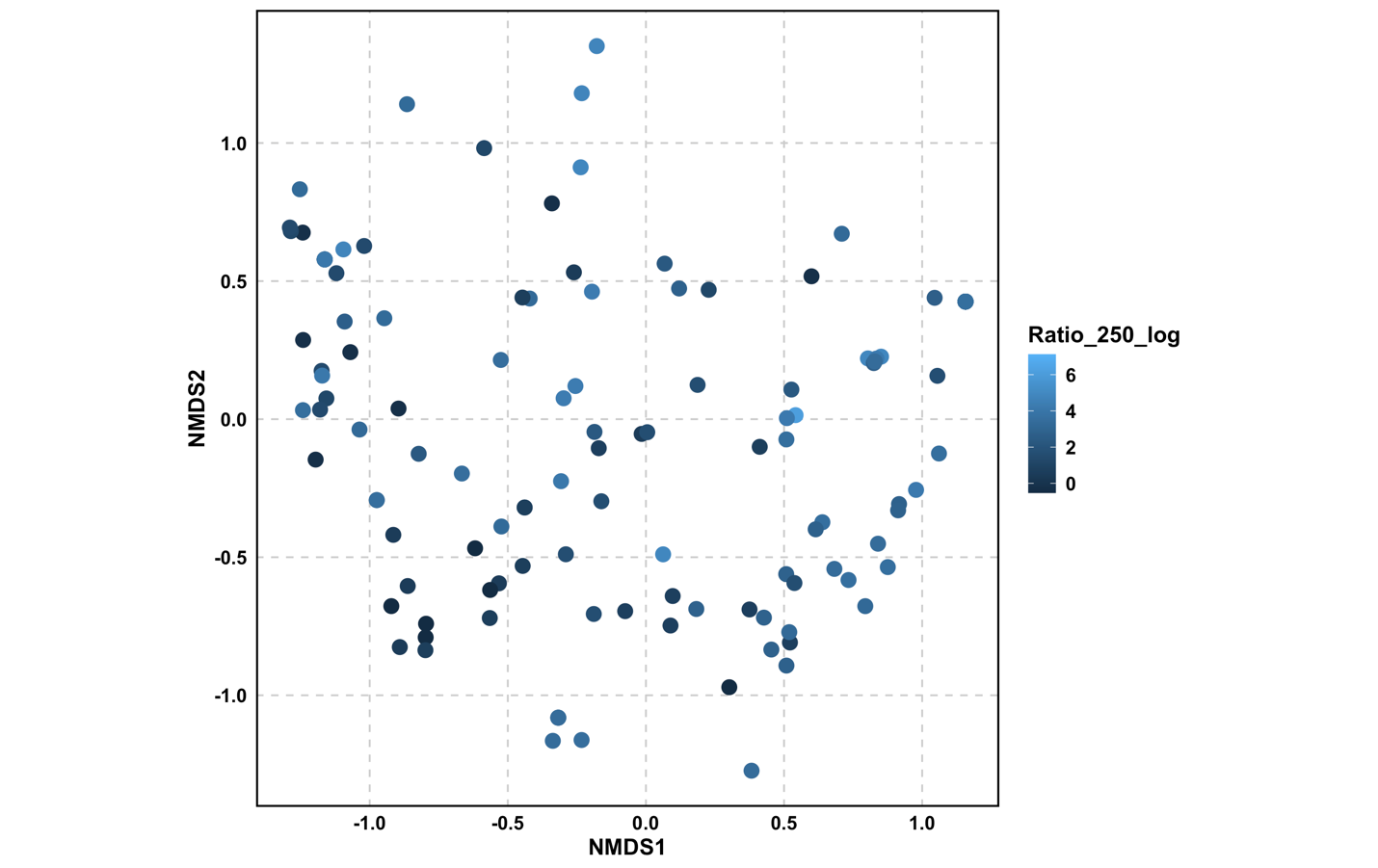

**Figure S7.** NMDS visualization of PERMANOVA of pollinator visitation to *Trifolium fucatum* by the log of the invasive to native plant ratio at 250 m.
